# Scouring the human Hsp70 network uncovers diverse chaperone safeguards buffering TDP-43 toxicity

**DOI:** 10.1101/2025.05.10.653282

**Authors:** Edward M. Barbieri, Miriam Linsenmeier, Katherine R. Whiteman, Yan Cheng, Sylvanne Braganza, Katie E. Copley, Paola Miranda-Castrodad, Brennen Lewis, Kevin Villafañe, Defne A. Amado, Beverly L. Davidson, James Shorter

## Abstract

Cytoplasmic aggregation and concomitant dysfunction of the prion-like, RNA-binding protein TDP-43 underpin several fatal neurodegenerative diseases, including amyotrophic lateral sclerosis. To elucidate endogenous defenses, we systematically scoured the entire human Hsp70 network for buffers of TDP-43 toxicity. We identify 30 J-domain proteins (2 DNAJAs, 10 DNAJBs, 18 DNAJCs), 6 Hsp70s, and 5 nucleotide-exchange factors that mitigate TDP-43 toxicity. Specific chaperones reduce TDP-43 aggregate burden and detoxify diverse synthetic or disease-linked TDP-43 variants. Sequence–activity mapping unveiled unexpected, modular mechanisms of chaperone-mediated protection. Typically, DNAJBs collaborate with Hsp70 to suppress TDP-43 toxicity, whereas DNAJCs act independently. In human cells, specific chaperones increase TDP-43 solubility and enhance viability under proteotoxic stress. Strikingly, spliceosome-associated DNAJC8 and DNAJC17 retain TDP-43 in the nucleus and promote liquid-phase behavior. Thus, we disambiguate a diverse chaperone arsenal embedded in the human proteostasis network that counters TDP-43 toxicity and illuminate mechanistic gateways for therapeutic intervention in TDP-43 proteinopathies.

---

Aberrant protein aggregation underlies several devastating neurodegenerative diseases such as amyotrophic lateral sclerosis (ALS) and frontotemporal dementia (FTD), for which there are currently no cures^1–3^. In healthy neurons, protein aggregation is opposed by intricate networks of molecular chaperones^4^, but these systems fail in the degenerating neurons of ALS/FTD patients. TDP-43, a primarily nuclear RNA-binding protein with a prion-like domain (PrLD)^5^, aggregates in the cytoplasm of afflicted neurons in ∼97% of ALS cases and ∼45% of FTD cases^6^. TDP-43 proteinopathy is also a feature of degenerating neurons in ∼57% of Alzheimer’s disease cases^7, 8^, all limbic-predominant age-related TDP-43 encephalopathy cases^9^, and ∼85% of chronic traumatic encephalopathy cases^10, 11^. TDP-43 has many functions in RNA metabolism, including pre-mRNA splicing and repression of cryptic exons^12, 13^. The propensity of TDP-43 to undergo phase transitions is critical for association with multiple nuclear biomolecular condensates, which enable maximal TDP-43 functionality^14, 15^. Most TDP-43 condensates in the nucleus contain RNA, which promotes their fluid-like properties^16^. However, under conditions of stress, TDP-43 can form nuclear bodies that are depleted of RNA and are more solid-like^17, 18^. Another important role for TDP-43 is to engage the NEAT1 long noncoding RNA (lncRNA) to regulate paraspeckles^19^, subnuclear membraneless organelles that modulate gene expression^20^. Dysregulation of paraspeckles and other nuclear condensates is associated with ALS/FTD^21^. Thus, pinpointing human molecular chaperones that antagonize aberrant TDP-43 phase transitions in the cytoplasm and the nucleus could provide avenues for therapeutic intervention to halt progression of ALS/FTD^5, 22, 23^.

The human genome encodes 194 known chaperones that can exhibit differential expression in different tissues, cell types, and with age^4, 24^. Within the vast proteostasis network^25^, the Hsp70 chaperone system comprised of J-domain proteins (JDPs), Hsp70 chaperones, and nucleotide-exchange factors (NEFs) regulate numerous housekeeping and stress-induced functions^26^. Indeed, mutations in components of the Hsp70 chaperone network are associated with disease^26^. Hsp70, consisting of a nucleotide-binding domain (NBD) linked to a substrate-binding domain (SBD), opposes protein aggregation and promotes protein folding through coordinated cycles of binding and release of client proteins^27^. These chaperone cycles are driven by ATP-dependent conformational changes in Hsp70 that are regulated by obligate co-chaperones, including JDPs and NEFs^26^. JDPs engage misfolded proteins and promote their transfer to Hsp70 via stimulation of ATP hydrolysis by Hsp70^26^. NEFs promote release of the client from Hsp70 by catalyzing exchange of ADP for ATP to restart the cycle^26^. Through iterative applications and variations of this basic cycle, the Hsp70 chaperone system maintains solubility and functionality of the entire proteome^28^.

JDPs contain a highly conserved ∼70 amino acid J-domain consisting of four helices that dock the JDP to Hsp70^29^. Functional J-domains contain a conserved histidine-proline-aspartate (HPD) tripeptide motif between helices II and III that is essential for stimulating Hsp70 ATPase activity^26^. JDPs are grouped into three classes based on common structural features. Class A and B JDPs share an N-terminal J-domain followed by a glycine-phenylalanine (GF)-rich linker and two ß-sandwich domains^26^. Class A JDPs also have a zinc finger-like region^26^. Class C JDPs share only the J-domain in common with other JDPs, and are otherwise highly diverse with a vast repertoire of domains that impart proteostasis in specialized contexts^26, 30^. For example, the Class C JDPs, DNAJC8 and DNAJC17 contain nuclear localization signals (NLSs) and long coiled-coil domains, enabling association with spliceosome components in the nucleus^31^. DNAJC17 contains an RNA-recognition motif (RRM) and localizes to nuclear speckles where it affects pre-mRNA splicing^32, 33^. However, in these and many other cases Class C JDPs remain enigmatic and poorly characterized.

In addition to their roles in the canonical Hsp70 chaperone cycle, JDPs and NEFs can function independently of Hsp70^34, 35^. For example, DNAJB6 and DNAJB8 antagonize polyglutamine aggregation without requiring a functional J-domain^36^, and Hsp110s prevent aggregation of misfolded proteins through a NEF-independent client-holding activity^37, 38^. Direct roles for JDPs, Hsp70s, and NEFs in buffering against TDP-43 aggregation and toxicity remain largely undetermined.

Here, we harnessed a powerful yeast model of TDP-43 proteinopathy^39, 40^ to systematically uncover human Hsp70-network chaperones that neutralize TDP-43 toxicity. Expression of human TDP-43 in yeast faithfully recapitulates key pathological features observed in degenerating neurons of TDP-43 proteinopathy patients, including cytoplasmic mislocalization, aggregation, and cell death^39, 40^. This model has served as a robust platform for discovering genetic and small-molecule modifiers of TDP-43 toxicity that have been validated in fly, mouse, human cells, and neuronal models of ALS/FTD^41–47^. Through this approach, we identified 41 human chaperones that suppress TDP-43 toxicity. Strikingly, specific chaperones reduce TDP-43 aggregation and buffer toxicity across both synthetic and disease-linked TDP-43 variants. We also uncovered minimal chaperone elements and unexpected, non-canonical mechanisms that counter TDP-43 toxicity. In human cells, these chaperones reduce insoluble TDP-43, enhance viability during proteotoxic stress, and promote nuclear TDP-43 localization. Collectively, our findings reveal a rich landscape of proteostatic defenses capable of restraining TDP-43 pathology and open new avenues for precision therapeutics targeting TDP-43-driven neurodegeneration.

## Results

### Multiple components of the human Hsp70 chaperone system mitigate TDP-43 toxicity

To study the entire human Hsp70 chaperone system in yeast we constructed a plasmid library encoding all 49 JDP genes plus 5 splice isoforms, all 12 Hsp70s, and all 14 NEF genes plus 2 splice isoforms under control of the galactose-inducible pGAL1 promoter. This library contains chaperones known to associate with a variety of subcellular compartments (Table S1, Figure S1A), enabling the search for TDP-43 safeguards throughout the cell. We tested the chaperones in two strains, expressing either a nontoxic mNeon protein^48^ or the toxic human TDP-43 (Figure 1A-E). The mNeon strain was used to assess growth effects caused by the chaperone independent of TDP-43. Compared to the vector control, there were no observed growth effects from human Hsp70s (Figure 1B, D) and only one NEF (BAG4) mildly impaired growth with mNeon (Figure 1C, D). By contrast, nearly half of the JDPs (26/54) exhibited at least a 10% growth defect (Figure 1A, D). Moreover, ten JDPs reduced growth more than 25% and two JDPs (DNAJB7 and DNAJC11) reduced growth below 50% of the vector control (Figure 1A, D). These results reveal that many human JDPs impact yeast growth even in the absence of TDP-43, highlighting the potential for broad cellular effects upon ectopic expression of JDPs. Moreover, this baseline analysis provides a critical reference for identifying specific components of the human Hsp70 network that modulate TDP-43 toxicity, independent of their intrinsic effects on cellular fitness.

**Figure 1.**
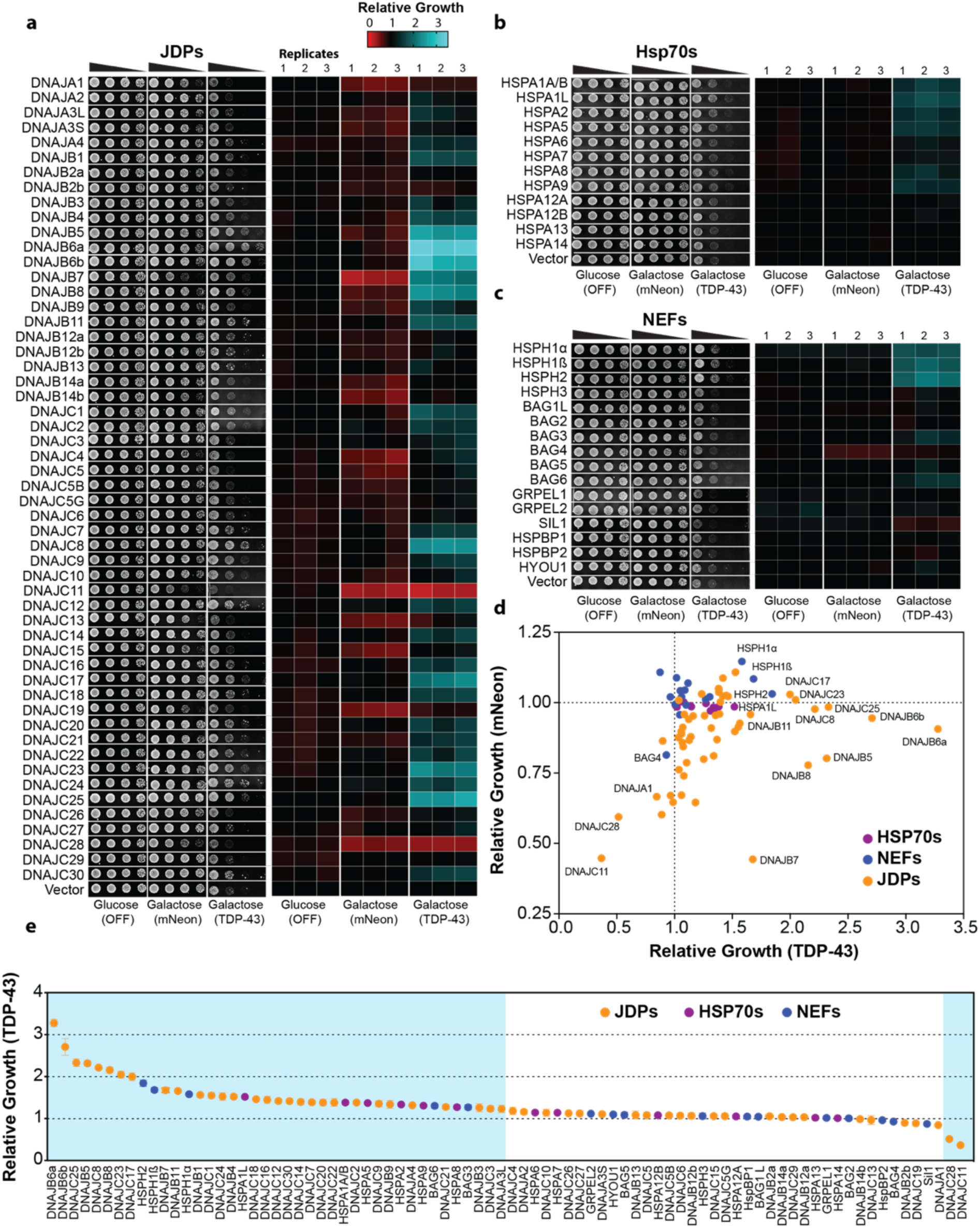
A yeast screen of human JDPs, Hsp70s, and NEFs identifies 41 components that suppress TDP-43 toxicity. (**A-C**) Yeast strains harboring galactose-inducible mNeon or TDP-43 were transformed with plasmids encoding galactose-inducible JDPs (A), Hsp70s (B), or NEFs (C). When grown on glucose media, there is no expression of mNeon, TDP-43, or chaperones. Strains were normalized to equivalent density (OD_600_ = 2), serially diluted 5-, 25-, and 125-fold, and the four cell densities were spotted onto glucose and galactose agar plates. Images show representative yeast growth alongside heatmap quantification from three independent replicates. Growth is shown relative to the vector control for each strain. **(D)** Combined 2D plot comparing relative growth of chaperones expressed in mNeon versus TDP-43 strains. Data points represent mean values from three replicates. **(E)** Rank-ordered plot of chaperones from the TDP-43 screen. Data are mean ± SEM for three replicates. Chaperones with a statistically significant difference from vector control (p < 0.05, one-way ANOVA and Dunnett’s multiple comparisons test) are highlighted in the blue box.

Elevated expression of TDP-43 poses a stringent test for the proteostasis network^39, 40^. Indeed, elevated expression of TDP-43 is linked to FTD^49^, and cytoplasmic TDP-43 aggregation is proposed to increase TDP-43 expression due to loss of TDP-43 autoregulation in a vicious cycle of disease^50, 51^. Remarkably, we uncovered 41 individual components of the human Hsp70 chaperone system that mitigate TDP-43 toxicity (Figure 1A-E). Thus, 50% of the diverse human Hsp70 chaperone system mitigate TDP-43 toxicity, indicating that this system provides a formidable proteostatic barrier that antagonizes aberrant TDP-43 behavior. These findings help explain why juvenile forms of ALS/FTD are exceedingly rare and that decline of the Hsp70 chaperone system during aging^52–55^ may increase the risk of developing TDP-43 proteinopathy. Moreover, the ability of diverse components of the human Hsp70 network to suppress TDP-43 toxicity in yeast suggests that boosting specific nodes of the Hsp70 chaperone network could have therapeutic benefit for TDP-43-related neurodegeneration.

### Nuclear, endoplasmic reticulum, and cytoplasmic components of the human Hsp70 network are protective against TDP-43

The set human Hsp70 network components that elicited toxicity in the mNeon strain was overrepresented (∼166%) in mitochondrial localization, suggesting that human chaperones can disrupt yeast mitochondrial homeostasis to impart toxicity (Figure S1B, C). Nuclear, cytoplasmic, and endoplasmic reticulum (ER)-associated chaperones were depleted in the toxic set by ∼40%, ∼11%, and ∼4% respectively. Among the 41 components that mitigate TDP-43 toxicity, ER and nuclear localization were enriched by ∼26% and ∼20% respectively, whereas mitochondrial chaperones were underrepresented by ∼33% (Figure S1D, E). The strongest suppressors of TDP-43 toxicity (>50% growth improvement) were enriched most highly for nuclear localization (∼32%) followed by ER (∼20%) and cytoplasm (∼6%) localization (Figure S1F, G). No mitochondrial chaperones were strongly protective against TDP-43. Therefore, mitigation of TDP-43 toxicity is most effective by chaperones known to localize to the nucleus, ER, and cytoplasm.

### Class C JDPs frequently antagonize TDP-43 toxicity

The majority of the 41 leads were JDPs (30/41), followed by Hsp70s (6/41), and NEFs (5/41; Figure 1E). Among the JDPs, Class A was generally ineffective, except for DNAJA4 and DNAJA3L, which mildly mitigated TDP-43 toxicity (Figure 1A, E). By contrast, 10 Class B and 18 Class C JDPs mitigated TDP-43 toxicity (Figure 1A, E). Thus, we uncover an unexpected proclivity for Class C JDPs, which encompass ∼44% of all leads, to antagonize TDP-43 toxicity. During evolution, Class C JDPs have undergone a remarkable adaptive radiation in multicellular organisms^56^. We suggest that the Class C JDP expansion may reflect, at least in part, a need to effectively buffer an increasing number of difficult, intrinsically aggregation-prone proteins like TDP-43^57^. Among these Class C JDPs was DNAJC7, which mildly mitigated TDP-43 toxicity (Figure 1A, E). Loss of function mutations in DNAJC7 are an established cause of ALS^58, 59^. Thus, loss of DNAJC7 function likely renders neurons more vulnerable to aberrant TDP-43 aggregation and toxicity, which drives ALS pathogenesis.

Our screen also identified known modifiers of aberrant TDP-43 behavior including DNAJB1^60^, DNAJB4^61^, DNAJB5^62^, DNAJB6^63–66^, HSPA1A^18^, HSPA1L^18^, HSPA5^18, 67^, HSPA8^18^, and BAG3^68^ (Figure 1A-E). Notably, we observed strong (>50%) suppression of TDP-43 toxicity from chaperones without established associations to TDP-43, including DNAJB7, DNAJB8, DNAJB11, DNAJC8, DNAJC17, DNAJC23, DNAJC24, and DNAJC25 (Figure 1A, E). DNAJB7 was unique in that it was toxic with mNeon, yet it suppressed TDP-43 growth defects strongly (Figure 1D). We also identified chaperones with mild (<50%) suppression of TDP-43 toxicity, including DNAJA3L, DNAJA4, DNAJB3, DNAJB9, DNAJC1, DNAJC2, DNAJC3, DNAJC9, DNAJC12, DNAJC14, DNAJC16, DNAJC18, DNAJC20, DNAJC21, DNAJC22, DNAJC30, HSPA2, HSPA9, and BAG6 (Figure 1A-E). Overall, our screen correctly identified known modifiers of aberrant TDP-43 behavior and revealed several new leads.

### Human Hsp70 network components that suppress TDP-43 toxicity typically modify TDP-43 aggregate number or size

TDP-43 normally shuttles between the nucleus and cytoplasm in human cells^69^. To evaluate how components of the human Hsp70 network that mitigate TDP-43 toxicity influence TDP-43 aggregation, we selected a panel of nuclear and cytoplasmic components for further analysis. These included seven JDPs (DNAJB5, DNAJB6a, DNAJB6b, DNAJB7, DNAJB8, DNAJC8, DNAJC17), an Hsp70 (HSPA1L), and three NEFs (HSPH1α, HSPH1β, and HSPH2). Notably, this panel did not act via reduction in TDP-43 expression or by inducing a heat-shock response, as HSP26 levels were not strongly increased (Figure S1H). To visualize and quantify the effects of these chaperones on TDP-43 aggregation, we employed a yeast strain expressing TDP-43-YFP. As a negative control, we included DNAJB2a, a JDP that does not suppress TDP-43 toxicity (Figure 1A, E). Expression of TDP-43-YFP was less toxic to yeast than untagged TDP-43, and most of the tested Hsp70 network components, except DNAJB7 and the negative control DNAJB2a, improved yeast growth in the presence of TDP-43-YFP (Figure S2A,B). DNAJB7 either fails to mitigate YFP-tagged TDP-43 toxicity or the intrinsic toxicity of DNAJB7 (Figure 1A, D) outweighs any protective effect.

As expected, TDP-43-YFP formed multiple cytoplasmic foci per cell, which were absent when YFP was expressed alone^39^ (Figure 2A). Co-expression of human Hsp70 network components with TDP-43-YFP revealed four distinct aggregation phenotypes:

a. Decreased number of TDP-43-YFP foci with unchanged average area per focus, observed with DNAJB5, DNAJB7, and DNAJC17 (Figure 2B-D; teal).
b. Decreased number of TDP-43-YFP foci with increased average area per focus, observed with DNAJB6a, DNAJB6b, HSPH1β, and HSPH2 (Figure 2B-D; yellow).
c. Increased number of TDP-43-YFP foci with decreased average area per focus, observed with DNAJB8, DNAJC8, and HSPH1α (Figure 2B-D; maroon).
d. Increased number of TDP-43-YFP foci with unchanged average area per focus, observed with HSPA1L and DNAJB2a (Figure 2B-D; blue).

**Figure 2.**
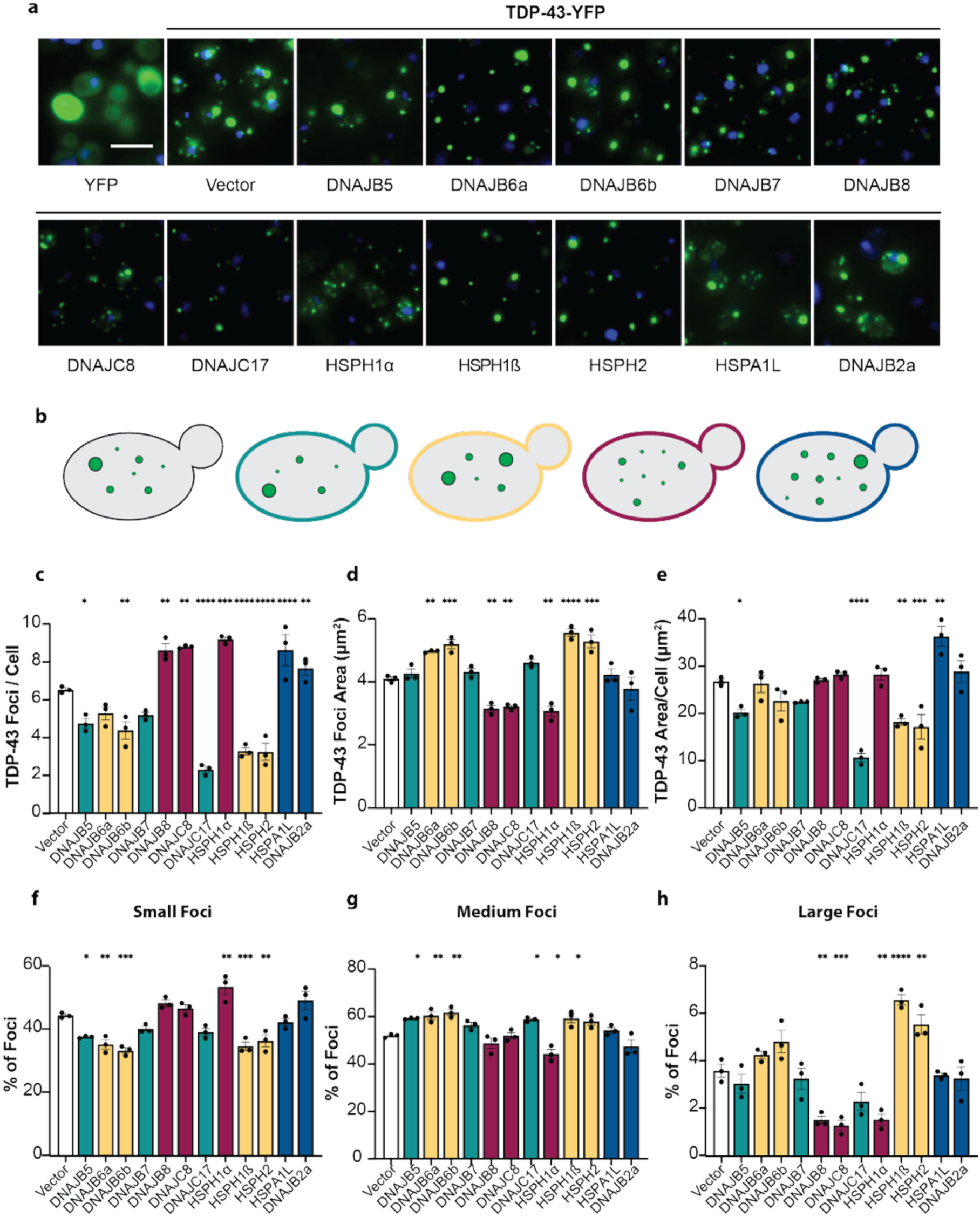
Components of the human Hsp70 network that suppress TDP-43 toxicity modify TDP-43 aggregate number or size. **(A)** Yeast expressing YFP alone (green) show diffuse signal throughout the cells, with DAPI staining nuclei (blue). Expression of TDP-43-YFP results in distinct cytoplasmic aggregates (green foci). Co-expression of TDP-43-YFP with the indicated human chaperones reveals diverse aggregate patterns compared to the empty vector control. Scale bar, 5 µm. **(B)** Schematic of yeast cells showing typical TDP-43-YFP aggregate burden for vector control (black outline) and four distinct aggregation phenotypes, color coded as follows: teal, decreased number of TDP-43-YFP foci, unchanged average area per focus; yellow, decreased number of TDP-43-YFP foci, increased average area per focus; maroon, increased number of TDP-43-YFP foci, decreased average area per focus; blue, increased number of TDP-43-YFP foci, unchanged average area per focus. **(C)** Number of TDP-43-YFP foci per cell. **(D)** TDP-43-YFP focus area. **(E)** Total TDP-43-YFP foci area per cell. **(F-H)** Percentage of small (F), medium (G), and large (H) foci. (C-H) Data are mean ± SEM from three independent replicates. Total number of cells analyzed: Vector, 2309; DNAJB5, 1631; DNAJB6a, 1995; DNAJB6b, 1595; DNAJB7, 4290; DNAJB8, 1718; DNAJC8, 758; DNAJC17, 1104; HSPH1α, 1247; HSPH1β, 1958; HSPH2, 2594; HSPA1L, 667; DNAJB2a, 245. (A-H) Statistical comparisons were made to the corresponding vector control using one-way ANOVA and Dunnett’s multiple comparisons test (*p < 0.05, **p < 0.01, ***p < 0.001, ****p < 0.0001).

Phenotypes (a) and (b) are consistent with previous studies indicating that a reduced number of TDP-43 foci per cell correlates with lower toxicity^39–41^. By contrast, an increased number of smaller or similarly sized TDP-43 foci as in phenotypes (c) and (d) has not been linked to protection against toxicity. Notably, only DNAJB5, DNAJC17, HSPH1β, and HSPH2 significantly reduced the total TDP-43 foci area, whereas only HSPA1L increased total foci area (Figure 2E). By classifying TDP-43 foci into three size categories (small (<1.3 µm²), medium (1.3-20.3 µm²), and large (>20.3 µm²)), we established that chaperones that reduced the number of TDP-43 foci typically decreased the proportion of small foci and increased the proportion of medium-sized foci (Figure 2C, F-H). Chaperones that increased the number of TDP-43 foci and decreased foci area (DNAJB8, DNAJC8, and HSPH1α) specifically decreased the proportion of large foci (Figure 2C-E, H). By contrast, HSPH1β and HSPH2 increased the proportion of large foci (Figure 2H). These observations point to multiple, mechanistically distinct modes by which Hsp70 network components can buffer TDP-43 proteotoxicity.

### Specific chaperones mitigate toxicity of disease-linked TDP-43 variants

Next, we tested the chaperones against a series of ALS/FTD-associated TDP-43 variants^6, 70–72^: P112H, K181E, G298S, Q331K, M337V, A382T, and I383V (Figure 3A, S2C-L). Typically, it is the wild-type version of TDP-43 that aggregates in disease, but in rare genetic forms of ALS/FTD (<1% of cases) a mutation in TDP-43 causes disease^72^. ALS/FTD-linked variants are typically found in the PrLD (e.g., G298S, Q331K, M337V, A382T, and I383V), but sometimes occur in RNA-recognition motif 1 (RRM1, as with P112H) or the linker between RRM1 and RRM2 (as with K181E)^72^. We found that the TDP-43 variants were expressed at similar levels (Figure S2L), but TDP-43^P112H^ was less toxic than TDP-43, whereas TDP-43^K181E^, TDP-43^G298S^, TDP-43^Q331K^, TDP-43^M337V^, TDP-43^A382T^, and TDP-43^I383V^ exhibited enhanced toxicity^40^ (Figure 3B).

**Figure 3.**
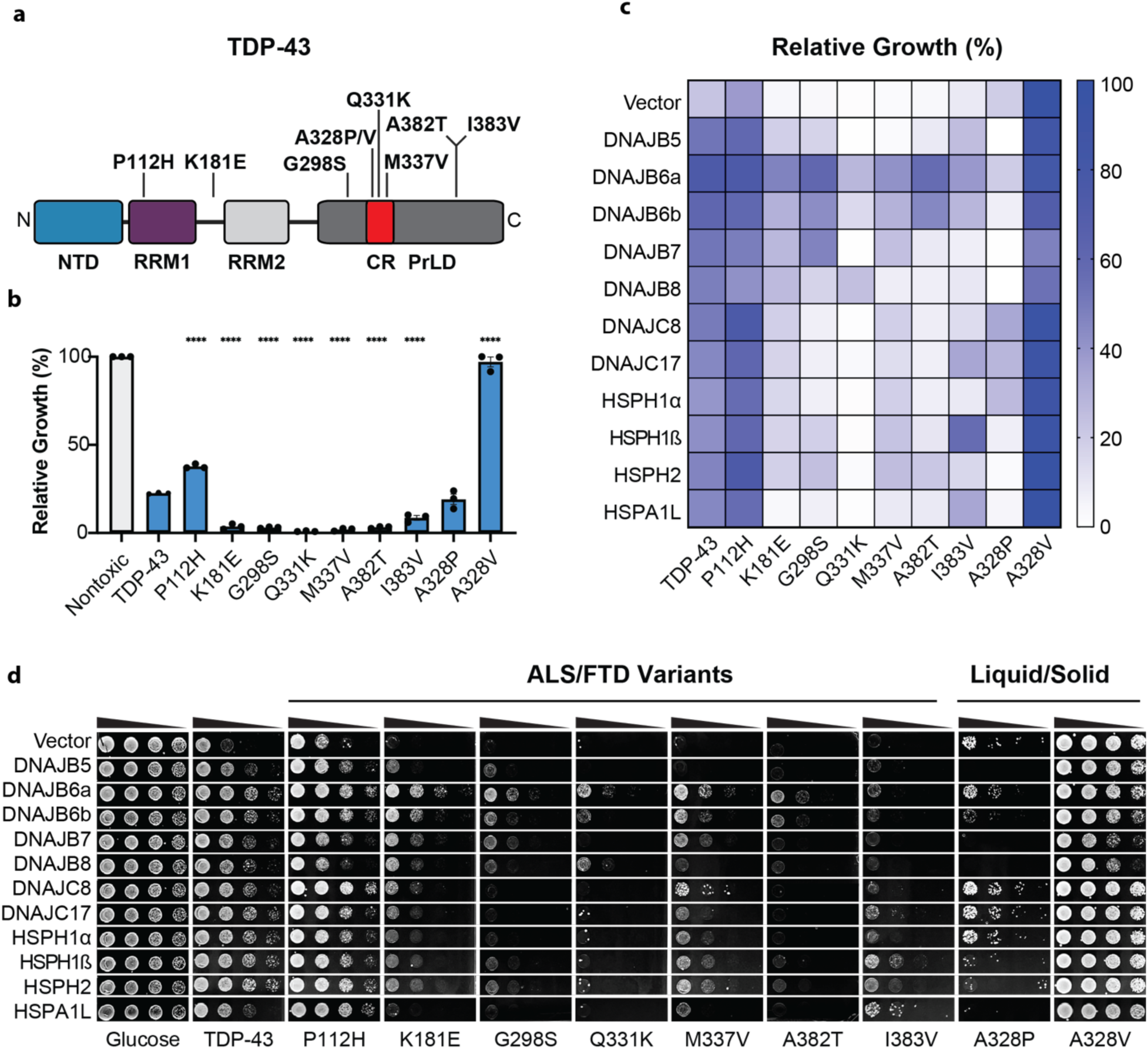
Chaperones have varied potency against disease-linked and synthetic TDP-43 variants. **(A)** Domain map of human TDP-43 showing the N-terminal domain (NTD), RNA recognition motifs (RRM1 and RRM2), prion-like domain (PrLD), and the conserved region (CR) at positions 320–343. Mutants tested are indicated along the domain map. **(B)** Relative yeast growth for the indicated TDP-43 variants, normalized to a nontoxic mNeon strain. Data are mean ± SEM from three independent replicates. Statistical comparisons were made to wild-type TDP-43 using one-way ANOVA and Dunnett’s multiple comparisons test (****p < 0.0001). **(C)** Heatmap quantification showing mean growth values normalized to the nontoxic mNeon strain for three independent replicates. **(D)** Yeast strains harboring galactose-inducible TDP-43 variants were transformed with plasmids encoding galactose-inducible chaperones. Strains were grown on glucose (no TDP-43 or chaperone expression) or galactose (induced expression) plates. Cultures were normalized to equivalent density (OD_600_ = 2), serially diluted 5-, 25-, and 125-fold, and spotted onto glucose and galactose agar plates. Representative yeast growth images are shown.

P112H is within RRM1, and exhibits altered RNA binding^71^. For TDP-43^P112H^, our chaperone panel mitigated toxicity, except for DNAJB8 (Figure 3C, D; S2C). This finding suggests that disease-linked mutations can enable TDP-43 to escape selective JDP buffers, which may contribute to disease pathogenesis.

K181E is in the linker between RRM1 and RRM2 and is reported to have altered RNA binding^73^. For TDP-43^K181E^, our chaperone panel mitigated toxicity, except for HSPA1L, with DNAJB6a conferring the most protection (Figure 3C, D; S2D). Notably, HSPA1L was the only chaperone that increased the total TDP-43-YFP foci area (Figure 2E), suggesting a distinct mechanism of toxicity suppression. Since HSPA1L exhibited reduced efficacy against a number of disease-linked TDP-43 variants (Figure 3C, D), this distinct mechanism appears to be readily evaded.

TDP-43^G298S^ is up to four times as stable as wild-type TDP-43 in human cells^74^, likely contributing to pathogenicity. Only the Class B JDPs, HSPH1β, and HSPH2 protected against TDP-43^G298S^ (Figure 3C,D; S2E). TDP-43^Q331K^ and TDP-43^M337V^ are in the transient α-helix in the PrLD (residues 320-343), which is critical for phase separation and aggregation^15, 75–80^. Indeed, these TDP-43 variants exhibit accelerated aggregation^40^ and enhanced stability^74^. TDP-43^Q331K^ toxicity was particularly difficult to suppress. Only DNAJB6a, DNAJB6b, and DNAJB8 improved growth (Figure 3C, D; S2F). By contrast, the chaperone panel reduced TDP-43^M337V^ toxicity, except for DNAJB5, DNAJB8, and HSPA1L (Figure 3C, D; S2G). For TDP-43^A382T^, only DNAJB6a and DNAJB6b strongly mitigated toxicity, whereas HSPH1β and HSPH2 provided weak protection (Figure 3C, D; S2H). Finally, for TDP-43^I383V^, only DNAJB5, DNAJB6a, DNAJB6b, DNAJC17, HSPH1ß, and HSPA1L conferred protection (Figure 3C, D; S2I).

In summary, TDP-43^P112H^, TDP-43^K181E^, and TDP-43^M337V^ toxicity was broadly buffered by the chaperone panel. TDP-43^G298S^ and TDP-43^I383V^ toxicity were more difficult to antagonize with fewer effective chaperones, whereas TDP-43^A382T^ and TDP-43^Q331K^ toxicity were the most challenging to overcome. These results highlight how single mutations in TDP-43, particularly in the PrLD, can reshape chaperone efficacy, even among highly similar isoforms such as HSPH1α and HSPH1β, which are >95% identical. Only DNAJB6a and DNAJB6b were effective against all tested disease-linked TDP-43 variants, suggesting they may have broad utility in combating TDP-43 proteinopathies. By contrast, other JDPs, Hsp70s, and NEFs may require a more precision-based approach, as therapeutic outcomes could vary depending on the underlying TDP-43 mutation. For instance, DNAJC8 provides the strongest protection against TDP-43^P112H^, but is ineffective against TDP-43^Q331K^.

### Specific chaperones mitigate toxicity of a synthetic TDP-43 liquid variant

TDP-43 assemblies can adopt distinct material states, ranging from dynamic condensates to solid-like aggregates ^81^. To examine how these states influence chaperone efficacy, we tested our panel against synthetic TDP-43 variants engineered to promote either solid-like or liquid-like behavior^82^. First, we tested TDP-43^A328V^, which forms nontoxic solid foci in yeast^82^. For nontoxic TDP-43^A328V^, the Class B JDPs modestly reduced growth to similar levels as in the mNeon strain (Figure 1A), whereas Class C JDPs, NEFs, and HSPA1L had no effect (Figure 3B-D; S2J). By contrast, TDP-43^A328P^ confers increase propensity to form toxic liquid-like foci in yeast^82^ and exhibits selective vulnerability to chaperone action. Only DNAJC8, DNAJC17, and HSPH1α could mitigate TDP-43^A328P^ toxicity (Figure 3B-D; S2K), indicating that these chaperones possess a unique capacity to buffer toxicity of TDP-43 condensates^82^. These findings suggest that the material properties of aberrant TDP-43 assemblies can also dictate which specific chaperones antagonize toxicity.

### Human Hsp110s and Hsp70 can passively buffer TDP-43 toxicity

We next investigated how human Hsp110 NEFs (HSPH1α, HSPH1β, and HSPH2) and the Hsp70 family member HSPA1L mitigate TDP-43 toxicity. Human Hsp110s are homologous to Hsp70s and share a similar domain architecture: an N-terminal NBD, a C-terminal ß-sheet-containing SBDß domain, followed by an α-helical SBDα “lid” domain, and a disordered tail (Figure 4A). Cytoplasmic Hsp70s, such as HSPA1L, also contain a conserved EEVD motif at the C-terminal end, which is a hotspot for co-chaperone interaction^83^. To determine whether a functional ATPase domain is required for Hsp110 and Hsp70 mitigation of TDP-43 toxicity, we constructed nonfunctional NBD mutants. A G233D mutation in the yeast Hsp110 homologue, SSE1, impairs ATP binding, Hsp70 interaction, and NEF activity^84–86^. The equivalent mutation in the human Hsp110s, G232D, also ablates these functions^87^. For HSPA1L, we introduced a D12N mutation analogous to the D10N mutation in HSPA1A, which eliminates ATPase activity^88, 89^. Surprisingly, these inactive NBD variants of HSPH1α, HSPH1ß, HSPH2 and HSPA1L retained their ability to suppress TDP-43 toxicity (Figure 4B-J). Thus, HSPH1α, HSPH1β, and HSPH2 suppress toxicity independently of Hsp70 interaction and NEF activity. Likewise, HSPA1L does not require the canonical JDP–Hsp70–NEF ATPase cycle to protect against TDP-43 toxicity.

**Figure 4.**
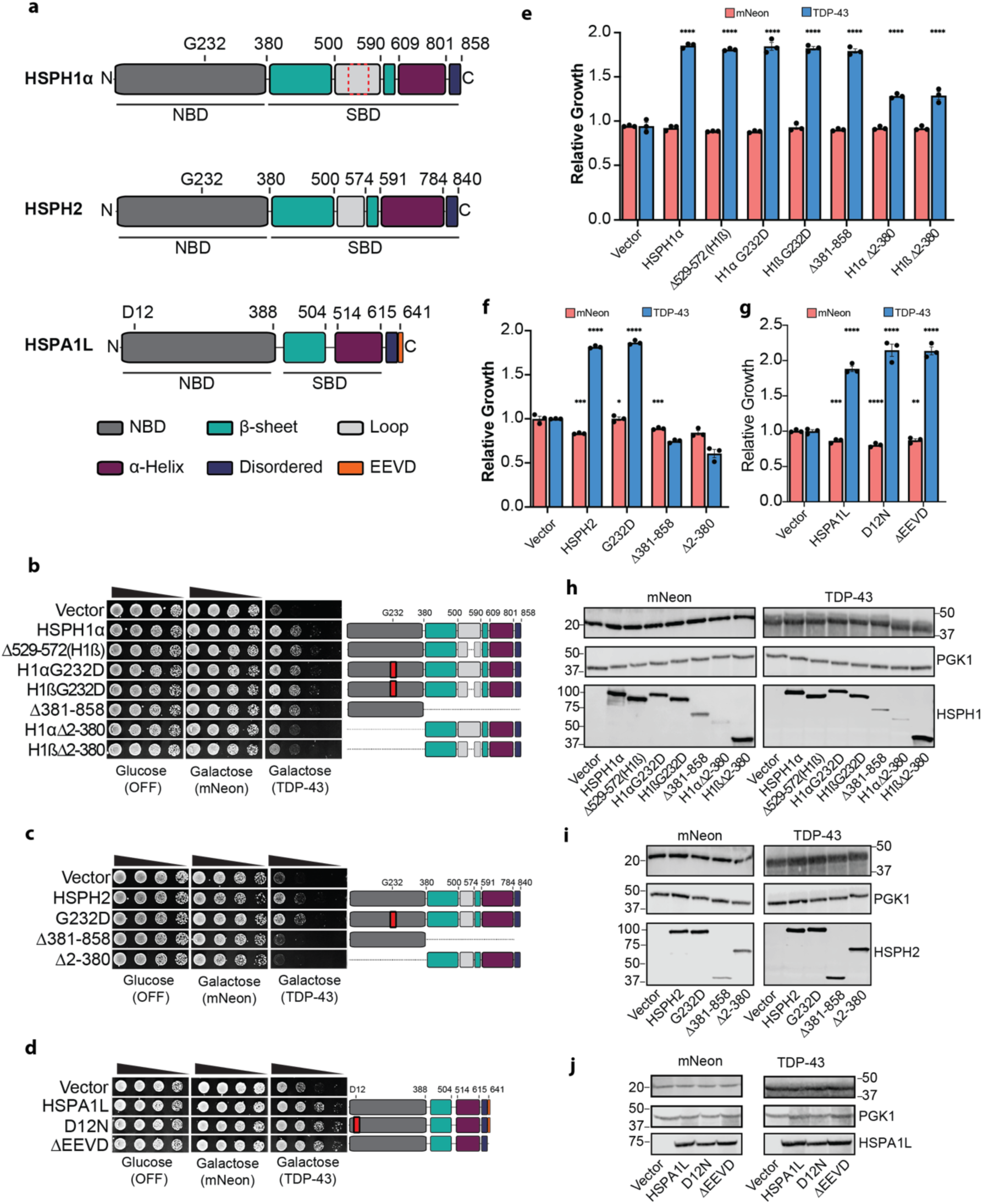
Human Hsp110s and Hsp70 passively buffer TDP-43 toxicity. **(A)** Domain maps of HSPH1α, HSPH2, and HSPA1L. HSPH1β, the HSPH1α Δ529–572 variant, is indicated by a dashed red outline. **(B-D)** Yeast strains harboring mNeon or TDP-43 were transformed with plasmids encoding the indicated mutants of HSPH1 (B), HSPH2 (C), and HSPA1L (D). Strains were grown on glucose (no mNeon, TDP-43, or chaperone expression) or galactose (induced expression) plates. Cultures were normalized to an equivalent density (OD_600_ = 2), serially diluted 5-, 25-, and 125-fold, and spotted onto glucose and galactose agar plates. Representative yeast growth images are shown. **(E-G)** Quantification of relative yeast growth in mNeon and TDP-43 strains for the indicated chaperone mutants. Values represent mean ± SEM from three independent replicates. Statistical comparisons were made to the corresponding vector control using one-way ANOVA and Dunnett’s multiple comparisons test (*p < 0.05, **p < 0.01, ***p < 0.001, ****p < 0.0001). **(H-J)** Corresponding Western blot images for HSPH1 (H), HSPH2 (I), and HSPA1L (J). PGK1 is used as a loading control; all chaperone variants are FLAG-tagged.

To further dissect the mechanism of Hsp110 action, we constructed deletion variants of HSPH1α, HSPH1β, and HSPH2 lacking either the N-terminal NBD or the C-terminal SBD (Figure 4A-D). HSPH1α and HSPH1ß are splice isoforms differing only in the SBD, where HSPH1ß lacks residues 529-572 (Figure 4A, dashed red box). HSPH1α is constitutively expressed and predominantly cytoplasmic, whereas HSPH1ß is expressed during mild heat shock and localized to the nucleus^90^. Intriguingly, the NBD of HSPH1α/ß was sufficient to mitigate TDP-43 toxicity to ∼80% of wild-type HSPH1α, despite being expressed at lower levels (Figure 4B, E, H). Moreover, the C-terminal SBD of HSPH1α or HSPH1ß partially suppressed TDP-43 toxicity to ∼50% of the full-length HSPH1 (Figure 4B, E, H). Thus, both the NBD and SBD of HSPH1α and HSPH1ß contribute to toxicity suppression, with the NBD playing a larger role. By contrast, neither the isolated NBD nor isolated SBD of HSPH2 could mitigate TDP-43 toxicity (Figure 4C, F, I). Rather, full-length HSPH2 was required to mitigate TDP-43 toxicity. Thus, HSPH1α, HSPH1β, and HSPH2 suppress TDP-43 toxicity through an NEF-independent mechanism but differ in domain requirements.

Tetratricopeptide repeat (TPR) domain proteins such as Hsp70/Hsp90 organizing protein (HOP) or C-terminus of Hsp70-interacting protein (CHIP) bind to the Hsp70 EEVD motif to act as adaptors to Hsp90 or the ubiquitin-proteasome system^91^. In addition, Class B JDPs interact with the Hsp70 EEVD motif to stimulate Hsp70 ATPase activity and client disaggregation^92^. Deletion of the EEVD motif did not impact HSPA1L mitigation of TDP-43 toxicity (Figure 4D, G, J). Overall, these unanticipated findings suggest that HSPH1α, HSPH1β, HSPH2, and HSPA1L can passively mitigate TDP-43 toxicity in a manner that bypasses traditional ATP-driven folding cycles and co-chaperone interactions.

### DNAJB5, DNAJB6a, DNAJB6b and DNAJB8 require either Hsp104 or Ssa1 to mitigate TDP-43 toxicity

To determine whether human chaperones act through conserved or yeast-specific mechanisms, we tested their ability to mitigate TDP-43 toxicity in strains lacking key yeast chaperones. Hsp104 is a protein disaggregase that uses ATP hydrolysis to dissolve protein aggregates^93, 94^, but has no direct homolog in humans^95^. This raises a key question: do human chaperones rely on yeast-specific machinery like Hsp104 to mitigate TDP-43 toxicity? To address this question, we first co-expressed the human chaperone panel with mNeon in a Δ*hsp104* strain. Toxicity profiles were largely unchanged compared to wild-type cells (Figure 1; S3A, B). Importantly, all chaperones except DNAJB7 continued to suppress TDP-43 toxicity in Δ*hsp104*, indicating that Hsp104 is not essential for their activity (Figure S3A, B).

Unlike Hsp104, Hsp70 is highly conserved from yeast to human^57^. The four major cytosolic Hsp70s in yeast, Ssa1-4, share high sequence identity with human Hsp70s^96^. Although deletion of all four SSA genes is lethal^97^, cells lacking three SSA genes (e.g., Δ*ssa2-4*) are viable^98^. We next tested the chaperone panel in a Δ*ssa1* strain. Similar to the Δ*hsp104* background, all chaperones except DNAJB7 suppressed TDP-43 toxicity (Figure S3C, D). These results suggest that most of the tested human chaperones suppress TDP-43 toxicity independently of Ssa1 alone.

Hsp104 and Ssa1 display a close functional relationship in yeast^99^. In the absence of Hsp104, Ssa1 can partially complement Hsp104 function and vice versa^99^. Thus, to explore potential redundancy between related Hsp104- and Ssa1-dependent pathways, we analyzed the human chaperone panel in the double-deletion strain Δ*hsp104*Δ*ssa1*. Strikingly, of the Class B JDPs, only DNAJB7 could mitigate TDP-43 toxicity in the Δ*hsp104*Δ*ssa1* strain (Figure S3E, F). This finding suggests that DNAJB5, DNAJB6a, DNAJB6b, and DNAJB8 require *either* Hsp104 *or* Ssa1 to mitigate TDP-43 toxicity.

DNAJB7 behaved differently than the other Class B JDPs. It was uniquely toxic in both Δ*hsp104* and Δ*ssa1* single deletions but not in the double-deletion background (Figure S3A-F). Thus, DNAJB7 may interact aberrantly with either Hsp104 or Ssa1 to elicit toxicity. In the absence of Hsp104 and Ssa1, this aberrant interaction is removed, allowing DNAJB7 to more effectively mitigate TDP-43 toxicity (Figure S3E, F).

In contrast, DNAJC8, DNAJC17, HSPA1L, HSPH1α, HSPH1β, and HSPH2 all suppressed TDP-43 toxicity in the Δ*hsp104Δssa1* strain (Figure S3E, F). These results highlight the ability of these chaperones to act independently of both Hsp104 and Ssa1. This autonomy also distinguishes these chaperones from Class B JDPs and underscores their broader functional flexibility.

To further investigate yeast SSA family contributions, we tested the human chaperone panel in the Δ*ssa2–4* strain (Figure S3G, H). This strain showed reduced fitness even when expressing mNeon, indicating heightened sensitivity (Figure S3G, H). Expression of DNAJB5 and HSPA1L exacerbated this toxicity (Figure S3G, H). Interestingly, HSPA1L could still mitigate TDP-43 toxicity in this strain, whereas DNAJB5 could not (Figure S3G, H). Thus, Δ*ssa2–4* cells may tolerate high HSPA1L levels more readily than high DNAJB5 levels under proteotoxic conditions of elevated TDP-43 concentrations. By contrast, DNAJB6a, DNAJB6b, and DNAJB8 suppressed TDP-43 toxicity in the Δ*ssa2–4* background, supporting their functional reliance on either Hsp104 or Ssa1 (Figure S3G, H). These results reinforce the concept that these Class B JDPs depend on at least one member of the Hsp104–Ssa1 axis.

DNAJB7 was not toxic in Δ*ssa2–4* cells (Figure S3G, H). However, DNAJB7 failed to mitigate TDP-43 toxicity in this background (Figure S3G, H). Thus, DNAJB7 may require one or more of Ssa2, Ssa3, or Ssa4 to confer protection against TDP-43 toxicity.

Finally, DNAJC8, DNAJC17, HSPH1α, HSPH1β, and HSPH2 remained protective in the Δ*ssa2– 4* background, further confirming that these chaperones do not rely on yeast Hsp104 or Ssa1-4. This persistent activity suggests that these chaperones act via pathways distinct from Hsp104 and Ssa1–4. Collectively, our findings uncover functional diversity within the human Hsp70 network and identify candidate chaperones most capable of overcoming cell-specific barriers to therapeutic translation.

### DNAJBs typically collaborate with Hsp70 to suppress TDP-43 toxicity, whereas DNAJCs act independently

To further dissect how Class B and Class C JDPs suppress TDP-43 toxicity, we focused on the role of the conserved ‘HPD’ motif within the J-domain, which is essential for stimulating Hsp70 ATPase activity^29^ (Figure 5A). Mutation of this motif to three alanines (HPD:AAA, hereafter mHPD) abolishes J-domain function^100, 101^. We thus generated mHPD variants across a panel of JDPs to test whether Hsp70 activation is required for suppression of TDP-43 toxicity (Figure 5B– D; S4A-M).

**Figure 5.**
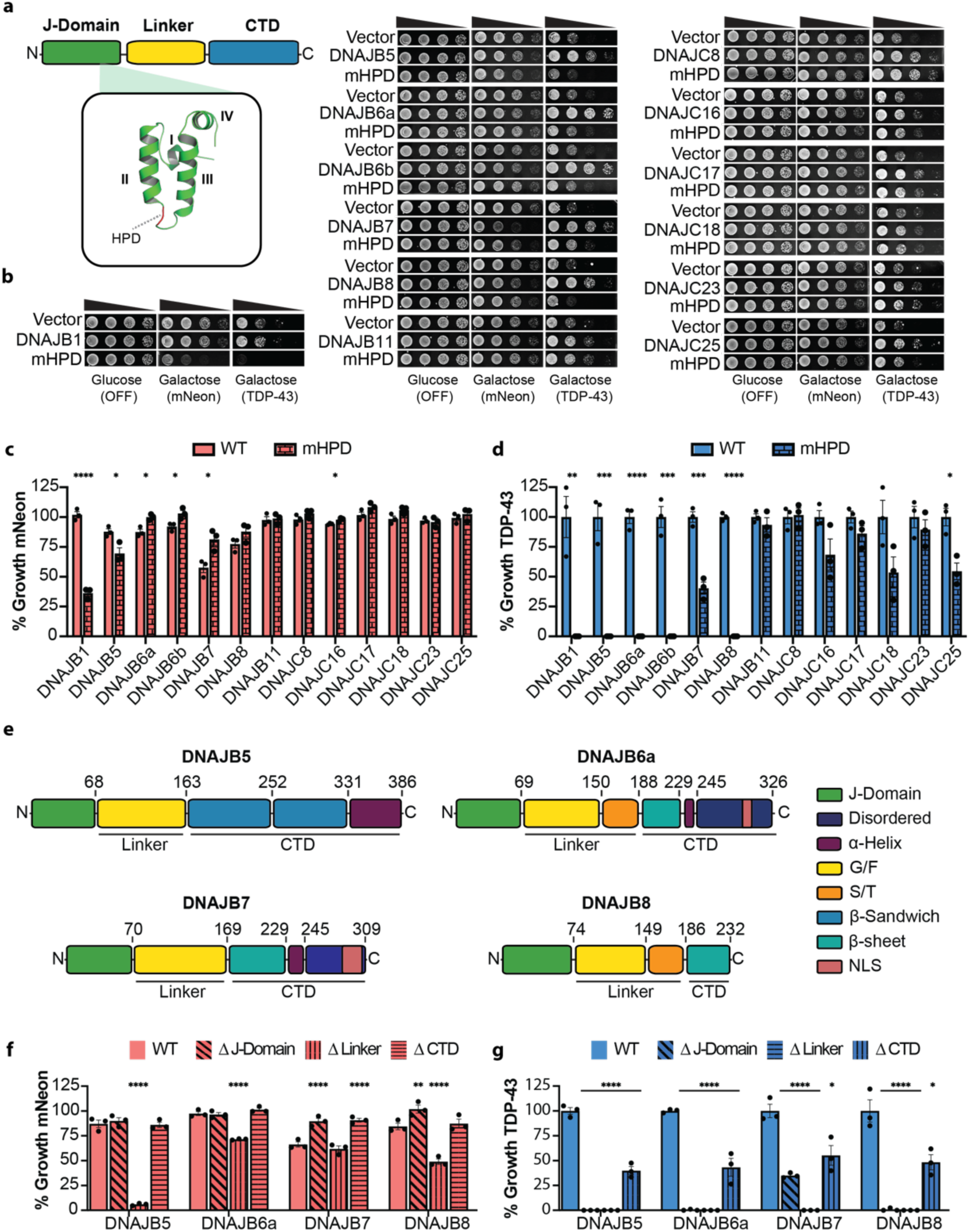
JDPs detoxify TDP-43 through Hsp70-dependent and Hsp70-independent mechanisms. **(A)** Representative domain map of a typical Class B JDP. The J-domain of human DNAJB1, as predicted by AlphaFold2^126^, is shown in green with helices I-IV indicated; the conserved HPD motif is shown in red. **(B)** Yeast strains harboring mNeon or TDP-43 were transformed with plasmids encoding the wild-type JDP or a corresponding HPD-to-AAA mutant (mHPD). Strains were grown on glucose (no mNeon, TDP-43, or chaperone expression) or galactose (induced expression) plates. Cultures were normalized to equivalent density (OD_600_ = 2), serially diluted 5-, 25-, and 125-fold, and spotted onto glucose and galactose plates. Representative growth images are shown for all JDPs tested. **(C)** Quantification of mNeon strain growth for wild-type JDPs and corresponding mHPDs. Growth is normalized to the nontoxic mNeon + vector strain. Values represent mean ± SEM from three independent replicates. **(D)** Quantification of TDP-43 strain growth for wild-type JDPs and corresponding mHPDs. For each mHPD, growth is normalized to the corresponding wild-type JDP. Values are mean ± SEM from three independent replicates. **(E)** Domain maps of DNAJB5, DNAJB6a, DNAJB7, and DNAJB8. **(F, G)** Quantification of domain deletion mutants for Class B JDPs in mNeon (F) and TDP-43 (G) strains. Values are mean ± SEM from three independent replicates. For (C,D), statistical significance was determined by a t-test between wild-type and corresponding mHPD (*p < 0.05, **p < 0.01, ***p < 0.001, ****p < 0.0001). For (F,G), significance was determined by one-way ANOVA relative to the vector control and Dunnett’s multiple comparisons test (*p < 0.05, **p < 0.01, ****p < 0.0001).

We first assessed the toxicity of each mHPD mutant in the mNeon control strain. All mHPD variants were expressed (Figure S5A-M). However, several Class B JDPs with mHPDs showed altered toxicity compared to wild type (Figure 5B, C; S4A-G). Specifically, mHPDs in DNAJB1 and DNAJB5 were more toxic than wild type (Figure 5B, C; S4A, B), likely acting as dominant negatives that compete with the yeast Class B JDP Sis1 for Ssa1 (Hsp70), given their high sequence identity^60^. In contrast, mHPD variants of DNAJB6a, DNAJB6b, DNAJB7, and DNAJB8 were less toxic than wild type, suggesting reduced interference with yeast Hsp70 (Figure 5B, C; S4C–F). Class C JDPs with mHPDs, by comparison, showed no change in toxicity (Figure 5B, C; S4H-M).

When TDP-43 was expressed, the functional dependence on Hsp70 sharply diverged between Class B and Class C JDPs (Figure 5B, D; S4A-M). Nearly all Class B JDPs with mHPDs lost the ability to suppress TDP-43 toxicity (Figure 5B, D; S4A-G). Only DNAJB7 mHPD retained partial activity, and only DNAJB11 mHPD was as protective as the wild type (Figure 5B, D; S4E, G). Thus, most Class B JDPs, including DNAJB5, DNAJB6a, DNAJB6b, and DNAJB8, require Hsp70 to mitigate TDP-43 toxicity, consistent with our findings in the Δ*hsp104*Δ*ssa1* strain (Figure S3E, F).

In sharp contrast, all Class C JDPs with mHPDs retained at least ∼50% activity (Figure 5B, D; S4H–M). Indeed, DNAJC8 and DNAJC17 mHPD variants matched the wild-type proteins in ability to suppress TDP-43 toxicity (Figure 5B, D; S4H–M). These results align with our genetic data in Δ*ssa1* and Δ*ssa2–4* strains, reinforcing the conclusion that Class C JDPs act independently of the SSA1–4 Hsp70s (Figure S3).

In summary, nearly all Class B JDPs suppress TDP-43 toxicity via Hsp70-dependent mechanisms, whereas Class C JDPs retain partial or full activity when they are unable to activate Hsp70. Thus, we reveal a fundamental mechanistic distinction, which indicates that Class C JDPs may directly chaperone TDP-43, bypassing the canonical JDP–Hsp70 axis.

### J-domain and linker are sufficient for DNAJB5, DNAJB6a, DNAJB7, and DNAJB8 to mitigate TDP-43 toxicity

To dissect how regions outside the J-domain contribute to suppression of TDP-43 toxicity by class B JDPs, we analyzed domain deletions in DNAJB5, DNAJB6a, DNAJB7, and DNAJB8 (Figure 5E–G; S6, S7). Consistent with mHPD results, deletion of the J-domain in DNAJB7 and DNAJB8 reduced toxicity in the mNeon strain, supporting the idea that these JDPs cause toxicity via excessive activation of Hsp70 (Figure 5F). Deletion of the conserved linker, which normally restrains promiscuous Hsp70 activation^63, 92^, caused strong toxicity, despite low expression, across all four class B JDPs tested (Figure 5F; S7). Indeed, even low expression of the isolated DNAJB5 J-domain (residues 1–68) was toxic, whereas the mHPD version was not (Figure S6A, B; S7A). Similarly, deletion of helix V (residues 94–113) in the DNAJB5 linker impaired growth, consistent with findings in DNAJB1 where helix V regulates J-domain activity^92^ (Figure S6A, B). By contrast, deletion of the C-terminal domain (CTD) had no effect on growth for any of the four JDPs (Figure 5F).

When TDP-43 was expressed, these domain dependencies became more pronounced. J-domain deletion abolished suppression of TDP-43 toxicity in all tested class B JDPs except DNAJB7, which retained partial activity (Figure 5G). Linker deletion eliminated toxicity suppression in all four JDPs (Figure 5G). Notably, ΔCTD constructs, comprising only the J-domain and linker, retained substantial protective activity, suppressing TDP-43 toxicity to over 40% of wild-type levels (Figure 5G). These results suggest that TDP-43 toxicity stems from insufficient activation of Hsp70. However, excessive or unregulated Hsp70 activity, such as that caused by deletion of the linker, can itself be toxic in the absence of stress. Thus, we reveal a critical therapeutic window where Hsp70 is activated just enough to suppress TDP-43 toxicity but not hyperactivated to elicit toxicity. This therapeutic window can be accessed by expression of the J-domain plus linker of several DNAJBs, which suggests minimal constructs for potential adeno-associated virus (AAV) therapies for TDP-43 proteinopathies. These results also suggest that pharmacological strategies to directly activate Hsp70 within an appropriate therapeutic window may be viable^102–104^.

### CTD-I and a specific splice isoform are required for DNAJB5 to mitigate TDP-43 toxicity

DNAJB5 is a canonical Class B JDP with an N-terminal J-domain, a glycine/phenylalanine-rich linker, and two C-terminal β-sandwich domains (CTD-I and CTD-II; Figure 5E), which protects against TDP-43 pathology in mice^62^. To further define which domains are essential for mitigating TDP-43 toxicity, we analyzed a series of deletion mutants targeting the β-sandwich domains, the C-terminal α-helical region, and a splice isoform with an alternative C-terminal sequence (Figure S6A,B; S7A, B). Most DNAJB5 deletion constructs were moderately to highly expressed in yeast. However, constructs 1–68, 1–68mHPD, 69–163, and Δ164–386 were only faintly detected (Figure S7A). Some constructs, including Δ2–68 and Δ69–93, showed signs of proteolytic cleavage (Figure S7A). Deletion of CTD-I (Δ164–252) impaired the ability of DNAJB5 to suppress TDP-43 toxicity, whereas deletion of CTD-II (Δ253–331) had no effect (Figure S6A, B; S7A). Removal of the C-terminal α-helical region (Δ332–386) only moderately reduced mitigation of TDP-43 toxicity (Figure S6A, B; S7A). We also compared the 386-amino acid (386aa) isoform of DNAJB5 (AAH12115.1) with a shorter, 348-amino acid (348aa) splice variant (UniProtKB O75953-3), which replaces the final 38 residues with an alternative 5-amino acid sequence (Figure S6A, B; S7A, B). Notably, the 348aa isoform failed to mitigate TDP-43 toxicity and exhibited mild toxicity on its own in the mNeon strain (Figure S6A, B; S7A, B). Together, these results demonstrate that CTD-I is essential for DNAJB5-mediated suppression of TDP-43 toxicity, whereas CTD-II and the α-helical tail are largely dispensable in the 386aa isoform. Moreover, the alternative C-terminal sequence in the 348aa splice variant abolishes protective activity, indicating that TDP-43 can elude specific chaperone isoforms.

### The DNAJB6a CTD suppresses TDP-43 toxicity

DNAJB6a, a noncanonical Class B JDP, is a potent suppressor of protein aggregation^36, 105^. Dominant mutations in the linker region of DNAJB6a cause limb girdle muscular dystrophy type 1D (LGMDD1), leading to myofibrillar protein aggregation^63^. The domain architecture of DNAJB6a includes a serine/threonine-rich (S/T) region in the linker and a C-terminal domain (CTD) composed of a single β-sheet domain, a short helix, and a predicted disordered tail that harbors an NLS (Figure 5E). All DNAJB6a variants were robustly expressed in yeast (Figure S7C). Although full-length DNAJB6a requires a functional J-domain to mitigate TDP-43 toxicity (Figure 5B, D), the CTD alone (Δ2–188) was sufficient to suppress TDP-43 toxicity in the absence of both the J-domain and linker (Figure S6C, D; S7C). Thus, the CTD may directly engage TDP-43. The β-sheet domain contributes to toxicity suppression, as deletion of residues 189–245 reduced suppression of TDP-43 toxicity by ∼50% (Figure S6C, D). By contrast, addition of the linker (Δ2– 69) eliminated activity (Figure S6C, D), indicating that the linker imposes autoinhibition on the CTD. The disordered tail containing the NLS was not required for activity (Δ246–326) (Figure S6C, D). Supporting these conclusions, the splice variant DNAJB6b, which lacks residues 242– 326, and contains an alternative 10-residue C-terminal sequence also suppressed TDP-43 toxicity (Figure 5B). These results establish the CTD as a central determinant of DNAJB6a-mediated protection. While the J-domain is essential in the context of the full-length protein, the CTD alone retains significant activity. This autonomous function points to both J-domain-dependent and independent mechanisms of DNAJB6a action. The CTD thus represents a minimal protective module and a compelling scaffold for engineering synthetic therapeutics that counteract TDP-43 proteinopathies.

### Nuclear localization underlies DNAJB7 toxicity and is not required to buffer TDP-43 toxicity

DNAJB7 is a testis-specific JDP initially identified as a tumor-associated antigen due to its absence in healthy tissues^106, 107^. This absence may reflect the toxicity of DNAJB7 when ectopically expressed (Figure 1A). Structurally, DNAJB7 combines a canonical J-domain and linker with a noncanonical CTD resembling that of DNAJB6a (Figure 5E). Most DNAJB7 constructs were expressed at moderate to high levels in yeast, except Δ2-168, Δ287-307, Δ245-309, and Δ169-309, which were weakly expressed (Figure S7D). Deletion of the disordered C-terminal tail (Δ245– 309) markedly increased toxicity in the mNeon strain, suggesting that this tail limits promiscuous or aberrant interactions (Figure S6E, F; S7D). Conversely, deletion of the J-domain (Δ2–69), N-terminal portion of the linker (Δ70–120), both J-domain and linker (Δ2–168), the nuclear localization signal (NLS; Δ287–307), or the entire CTD (Δ169–309) reduced toxicity (Figure S6E,F; S7D). Strikingly, removal of the NLS nearly eliminated DNAJB7 toxicity, implicating nuclear localization as a key driver of harmful activity. Together with the mHPD data (Figure 5B,D), these results suggest that DNAJB7 promotes toxicity through inappropriate Hsp70 activation and Hsp70-independent mechanisms. In the absence of the regulatory C-terminal tail containing the NLS, additional toxicity may be unleashed in the cytoplasm.

In the TDP-43 strain, DNAJB7 domain deletions resembled those observed for DNAJB6a. Expression of the CTD alone (Δ2–168) was sufficient to suppress TDP-43 toxicity (Figure S6E, F). Deletion of the β-sheet domain (Δ169–229) lessened this protective effect, whereas removal of the adjacent α-helix (Δ230–244) had minimal impact (Figure S6E, F). Importantly, nuclear localization was dispensable, as deletion of the NLS (Δ287–307) preserved suppression of TDP-43 toxicity (Figure S6E, F). These findings establish the CTD of DNAJB7 as a minimal, functional module capable of mitigating TDP-43 toxicity. Moreover, because nuclear localization is not required for protection, redirecting DNAJB7 to the cytoplasm could minimize toxicity while preserving therapeutic activity.

### DNAJB8 oligomerization is not required to buffer TDP-43 toxicity

DNAJB8 is highly expressed in the testis but undetectable in the human brain^108^. In cell models, DNAJB8 suppresses aggregation of polyglutamine peptides^36^. This JDP shares 63% sequence identity with DNAJB6b and features a J-domain, a linker containing a serine/threonine-rich (S/T) region, and a C-terminal β-sheet domain (Figure 5E). DNAJB8 was highly expressed in yeast with some evidence of proteolysis, and the deletion constructs were expressed at lower levels than the wild-type DNAJB8 (Figure S7E). DNAJB8 can form oligomers through the S/T region, yet monomeric variants retain the ability to bind aggregation-prone substrates and suppress aggregation^109^. In the TDP-43 strain, deletion of the S/T region (Δ149–186) did not impair suppression of TDP-43 toxicity (Figure S6G, H; S7E). These findings indicate that oligomerization via the S/T region is not required for the protective activity of DNAJB8 against TDP-43 toxicity.

### The J-Domain and adjacent helix of DNAJC8 mitigate TDP-43 toxicity independently of Hsp70

DNAJC8 is a predominantly nuclear JDP that interacts with spliceosome components^31^ and directly associates with the splicing regulator SRPK1^110^. DNAJC8 suppresses polyglutamine aggregation in a J-domain-independent manner^111^. Structurally, DNAJC8 contains a J-domain flanked by short helices, a long coiled-coil domain with predicted NLSs, and a C-terminal disordered tail (Figure 6A). DNAJC8 deletion constructs were detected at low to moderate expression levels in yeast (Figure S8A). Although loss of canonical J-domain function (mHPD) had no impact on DNAJC8 toxicity in the mNeon strain (Figure 5B, C), deletion of the J-domain (Δ57–124) caused strong toxicity (Figure 6A-C; S8A). This toxicity was mapped to the coiled-coil domain (residues 153–231), as deletion of this region (Δ153–231) abolished toxicity, whereas deletion of residues 2–152 or 57–124 alone remained toxic. Notably, simultaneous deletion of the J-domain and the disordered C-terminal tail (Δ57–124Δ232–253) was the most toxic combination (Figure 6B, C). These findings suggest that the J-domain, and to a lesser extent, the disordered C-terminal tail, suppresses unwanted interactions involving the coiled-coil domain. Supporting this idea, deletion of the helix adjacent to the J-domain (Δ125–143) caused mild toxicity (Figure 6B, C), possibly by impairing regulation of the coiled-coil region.

**Figure 6.**
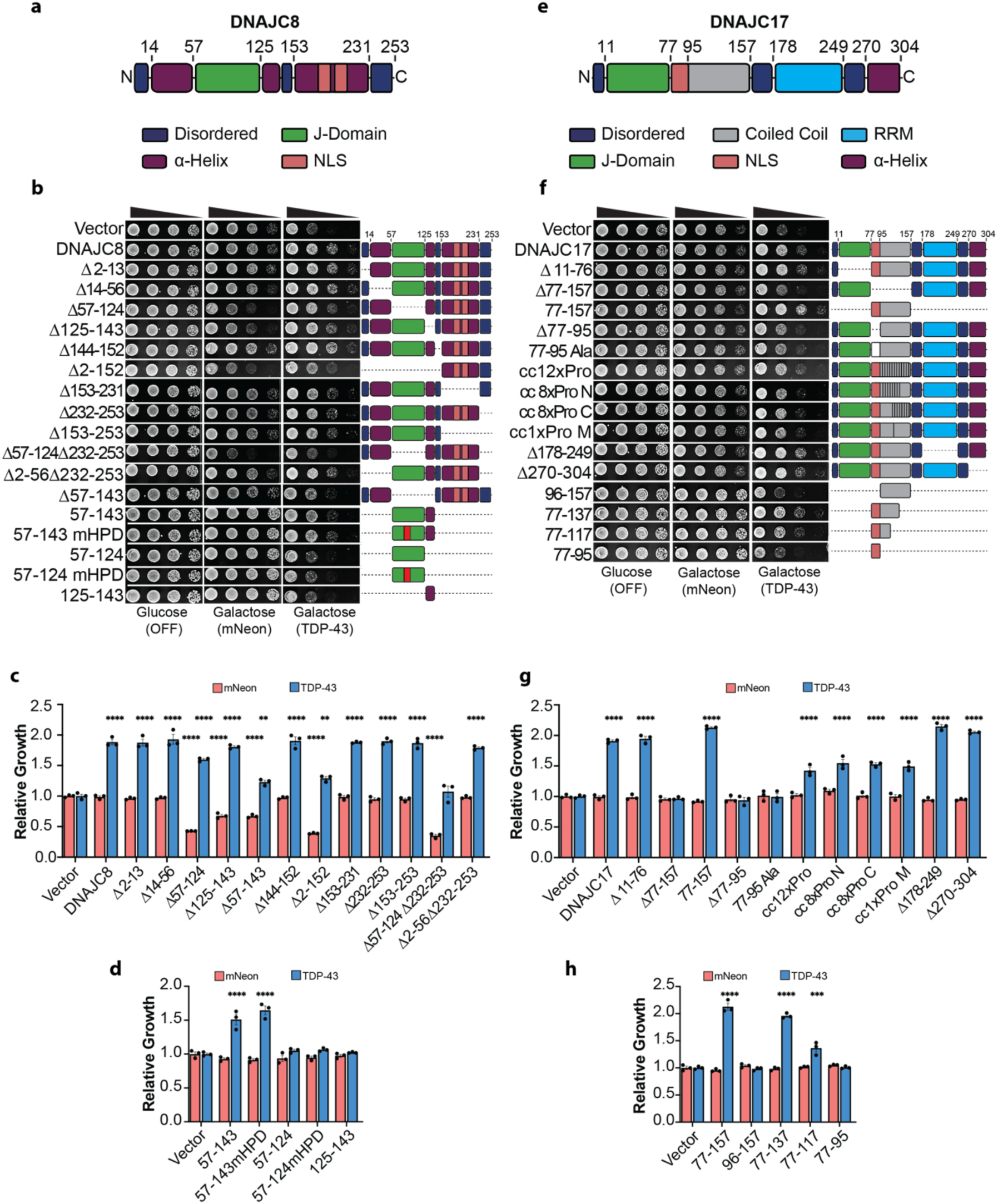
Domain mapping reveals minimal regions required for DNAJC8 and DNAJC17 suppression of TDP-43 toxicity. **(A)** Domain map of DNAJC8. **(B)** Yeast strains harboring mNeon or TDP-43 were transformed with plasmids encoding DNAJC8 domain deletion constructs. Strains were grown on glucose (no mNeon, TDP-43, or chaperone expression) or galactose (induced expression) plates. Cultures were normalized to equivalent density (OD_600_ = 2), serially diluted 5-, 25-, and 125-fold, and spotted onto glucose and galactose plates. **(C,D)** Quantification of yeast growth assays shown in (B). **(E)** Domain map of DNAJC17. **(F)** Representative yeast growth assays for DNAJC17 domain deletion constructs. **(G,H)** Quantification of yeast growth assays shown in (F).Values are mean ± SEM from three independent replicates. Statistical significance was determined relative to the vector control by one-way ANOVA and Dunnett’s multiple comparisons test (**p < 0.01, ***p < 0.001, ****p < 0.0001).

In the TDP-43 strain, most individual deletions, including residues 232–253, previously linked to polyglutamine suppression^111^, had little effect on TDP-43 toxicity mitigation (Figure 6B–D). These findings suggest that multiple domains collectively contribute to the DNAJC8 buffer, and loss of a single region is generally insufficient to abrogate protection. However, deletion of the J-domain (Δ57–124) or the adjacent helix (Δ125–143) modestly reduced suppression of TDP-43 toxicity, and loss of both (Δ57–143) further impaired protection (Figure 6B, C). Interestingly, deletion of the N-terminal region (Δ2–152), which was toxic in the mNeon strain, retained robust suppression of TDP-43 toxicity (Figure 6B, C), indicating that the C-terminal portion alone safeguards against TDP-43 toxicity.

Since canonical J-domain activity through Hsp70 stimulation is dispensable for suppression of TDP-43 toxicity by DNAJC8 (Figure 5B, D), yet the 57–143 region comprising the J-domain and adjacent helix is important to buffer TDP-43 toxicity, we hypothesized that this region reduces TDP-43 toxicity in an Hsp70-independent manner. Thus, we analyzed fragments corresponding to residues 57–143, a J-domain-inactive variant (57–143:mHPD), the J-domain alone (57–124), 57– 124:mHPD, and the adjacent helix (125–143) (Figure 6D). These fragments were not detected by Western blot (Figure S8A), likely due to low expression and small size. Nonetheless, the 57–143 fragment suppressed TDP-43 toxicity to ∼80% of full-length DNAJC8 levels, and 57–143:mHPD was surprisingly even more effective, reducing toxicity to ∼85% of wild-type levels (Figure 6D). In contrast, 57–124, 57–124:mHPD, and 125–143 were ineffective (Figure 6D). Together, these results suggest that multiple regions of DNAJC8 contribute to protection against TDP-43 toxicity, with the 57–143 region playing a central role. Enhanced suppression by the J-domain-inactive 57– 143:mHPD variant indicates that direct interaction between this region and TDP-43, rather than stimulation of Hsp70, may drive the protective effect. To our knowledge, direct engagement of a substrate by a J-domain to mitigate toxicity has not been described. Alternatively, the 57–143 fragment may modulate Hsp70 activity through a noncanonical mechanism to confer protection.

### DNAJC17 suppresses TDP-43 toxicity via the coiled-coil region and nuclear localization

DNAJC17 is an essential, predominantly nuclear JDP that interacts with spliceosome components^31^ and localizes to nuclear speckles^32^. The yeast ortholog, Cwc23, is required for spliceosome disassembly^112^, a function that is independent of the J-domain^113^. In human cells, DNAJC17 knockdown disrupts pre-mRNA splicing and causes exon skipping in genes involved in cell-cycle progression^33^. Notably, DNAJC17 is the only JDP that contains an RRM. DNAJC17 domain architecture includes an N-terminal J-domain, a long coiled-coil region (residues 77–157), an RRM, and a C-terminal helix (Figure 6E). All DNAJC17 deletion constructs were expressed and were nontoxic in the mNeon strain, with only 77-157 exhibiting low detection (Figure 6F–H; S8B, C).

In the TDP-43 strain, deletion of the J-domain (Δ11–76) or the C-terminal helix (Δ270–304) of DNAJC17 had no impact on TDP-43 toxicity suppression (Figure 6F–H). By contrast, removal of the coiled-coil region (Δ77–157) abolished the protective effect (Figure 6F, G). Unexpectedly, deletion of the RRM (Δ178–249) enhanced suppression of TDP-43 toxicity by DNJAC17 (Figure 6F, G). Since the RRM inhibits J-domain–mediated stimulation of Hsp70 ATPase activity^33^, this enhancement may reflect increased Hsp70 activation. Alternatively, loss of the RRM may reduce off-target interactions, increasing specificity toward TDP-43.

Protection against TDP-43 toxicity was lost upon deletion of the NLS (Δ77–95) of DNAJC17 or substitution of this region with alanines (77–95Ala), indicating that nuclear localization is essential (Figure 6F, G). To further assess the contribution of the coiled-coil region, proline insertions were introduced to disrupt α-helical structure. A single proline in the center of the coiled coil (cc1xPro M) reduced TDP-43 toxicity suppression, whereas constructs with eight prolines at either end (cc8xPro N,C) or 12 distributed prolines (cc12xPro) did not further impair activity (Figure 6F, G).

Expression of residues 77–157 alone strongly suppressed TDP-43 toxicity and slightly outperformed full-length DNAJC17 (Figure 6F, G). FLAG-tagged constructs confirmed this result (Figure 6F, H; S8C). Consistent with full-length DNAJC17 behavior, removal of the NLS (construct 96–157) eliminated activity (Figure 6F, H; S8C). C-terminal truncations defined a minimal protective fragment spanning residues 77–137, which was as effective as the full-length protein, while 77–117 retained partial activity (Figure 6F, H; S8C). The isolated NLS region (77– 95) was insufficient for suppression, although low expression of the 77–117 and 77–95 fragments may contribute to this insufficiency (Figure 6F, H; S8C). Together, these findings establish the coiled-coil region of DNAJC17 as both necessary and sufficient to mitigate TDP-43 toxicity. The requirement for nuclear localization further supports a mechanism involving nuclear compartmentalization, potentially through engagement with splicing-related factors.

### JDPs and HSPH1 isoforms reduce TDP-43 aggregation in human cells

To extend findings from the yeast model, we examined how DNAJB5, DNAJB6a, DNAJB6b, DNAJC8, DNAJC17, HSPH1α, and HSPH1β influence TDP-43 proteostasis in human cells. Human (HEK293) cells were co-transfected with V5-tagged chaperone constructs and either TDP-43-YFP or a TDP-43-YFP variant with an inactivated nuclear localization signal (TDP-43mNLS-YFP), and localization was assessed (Figure 7A; S9A). In contrast to YFP alone, which was diffusely distributed, TDP-43-YFP localized primarily to the nucleus under all conditions, regardless of chaperone expression (Figure 7A). As expected, TDP-43mNLS-YFP was confined to the cytoplasm (Figure S9A). The chaperones displayed distinct localization patterns: DNAJB5, DNAJB6b and both HSPH1 isoforms were distributed across nucleus and cytoplasm, whereas DNAJB6a, DNAJC8, and DNAJC17 localized to the nucleus, (Figure 7A; S9A). To quantify TDP-43 aggregation, we measured RIPA-insoluble TDP-43 by fractionating lysates and performing Western blot analysis. TDP-43-YFP expression increased insoluble TDP-43 by ∼49% relative to YFP alone, whereas TDP-43mNLS-YFP caused a larger increase of ∼170% (Figure 7B). DNAJB5, DNAJB6a, DNAJB6b, DNAJC17, HSPH1α, and HSPH1β reduced insoluble TDP-43 levels by ∼20% compared to the vector control, with DNAJC8 showing the strongest effect, lowering insoluble TDP-43 by ∼35% (Figure 7C). In the case of TDP-43mNLS-YFP, which was more resistant to chaperone activity, only DNAJB6a, DNAJB6b, and DNAJC17 reduced aggregation by ∼25%, whereas DNAJC8 again had the most potent effect, decreasing insoluble TDP-43 by ∼50% (Figure 7D). These findings demonstrate that several human JDPs and HSPH1 isoforms reduce TDP-43 aggregation in human cells, with DNAJC8 emerging as the most effective at lowering insoluble TDP-43 levels.

**Figure 7.**
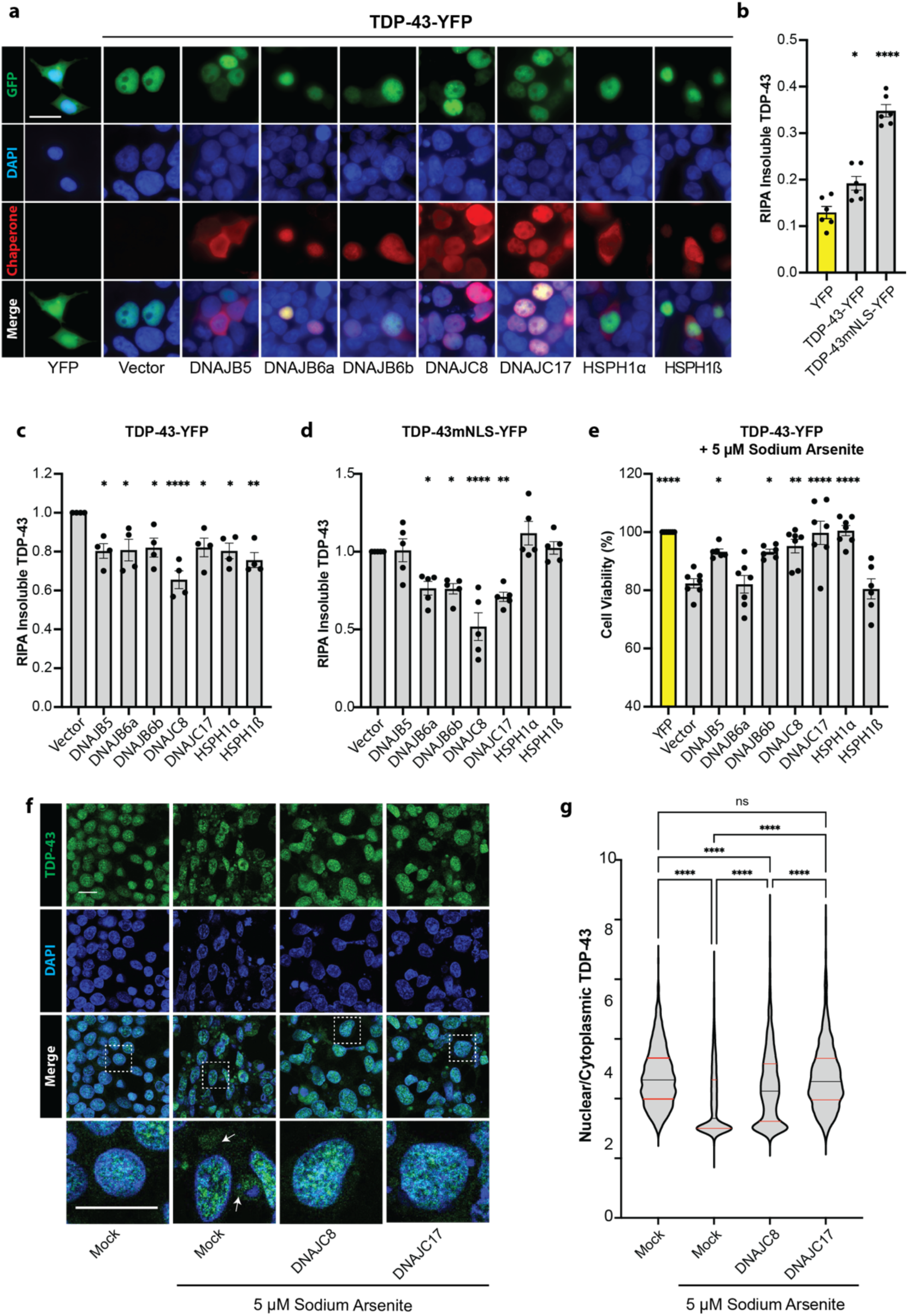
Chaperones safeguard against TDP-43 insolubility, mislocalization, and toxicity in human cells. **(A)** Representative images showing expression of YFP, TDP-43-YFP, and chaperones in HEK293 cells. Cell nuclei are stained with DAPI, and V5-tagged chaperones are detected by immunofluorescence. Scale bar, 20 µm. **(B)** Quantification of total RIPA-insoluble TDP-43 in cells transfected with YFP, TDP-43-YFP, or TDP-43mNLS-YFP. Values represent mean ± SEM from six replicates. Statistical significance was determined by one-way ANOVA relative to the YFP condition and Dunnett’s multiple comparisons test (*p < 0.05, ****p < 0.0001). **(C)** Quantification of RIPA-insoluble TDP-43 normalized to the vector control for TDP-43-YFP. Values represent mean ± SEM from four replicates. **(D)** Quantification of RIPA-insoluble TDP-43 normalized to the vector control for TDP-43mNLS-YFP. Values represent mean ± SEM from five replicates. **(E)** Cell viability in cells transfected with TDP-43-YFP and chaperones, followed by treatment with 5 µM sodium arsenite for 48 hours post-transfection. Values are normalized to the YFP control in each experiment and represent mean ± SEM from seven replicates. Statistical significance for (C–E) was determined by one-way ANOVA relative to the vector control and Dunnett’s multiple comparisons test (*p < 0.05, **p < 0.01, ****p < 0.0001). **(F)** Representative images from TDP-43 nuclear retention assay after 48-hour treatment with 5 µM sodium arsenite post-transfection. Nuclei are stained with DAPI, and endogenous TDP-43 is detected by immunofluorescence. Areas captured in magnified images are marked by dashed boxes. White arrows indicate examples of cytoplasmic TDP-43. Scale bars, 20 µm. **(G)** Quantification of the TDP-43 nuclear-to-cytoplasmic ratio. Number of cells quantified: n = 1771 (Mock untreated), n = 1345 (Mock), n = 1217 (DNAJC8), and n = 1065 (DNAJC17). Black lines represent the median; red lines represent the quartiles. Statistical significance was determined by one-way ANOVA and Tukey’s test ( ****p < 0.0001).

### Chaperones protect against TDP-43 toxicity under chronic stress

Expression of TDP-43-YFP or TDP-43mNLS-YFP caused only mild toxicity in HEK293 cells, reducing viability by ∼5% and ∼9%, respectively, compared to the YFP control (Figure S9B). However, under chronic low-dose stress (5 µM sodium arsenite for 48 hours), TDP-43-YFP further reduced viability by ∼20%, whereas TDP-43mNLS-YFP had no additional effect (Figure S9B). To evaluate chaperone protection under these conditions, we co-expressed chaperones with TDP-43-YFP and treated cells with sodium arsenite for 48 hours post-transfection. DNAJB5 and DNAJB6b improved viability by ∼13% relative to the vector control, DNAJC8 increased viability by ∼16%, and DNAJC17 and HSPH1α fully restored viability to the level of the YFP control (Figure 7E). These findings indicate that while TDP-43 expression alone is only mildly toxic in HEK293 cells, chronic stress enhances toxicity. Several chaperones, especially DNAJC17 and HSPH1α, provided strong protection, fully restoring cell viability and highlighting their potential to counteract stress-induced TDP-43 proteinopathy.

### DNAJC8 and DNAJC17 enhance TDP-43 nuclear retention under chronic stress

Chronic stress is known to induce the mislocalization of TDP-43 from the nucleus to the cytoplasm, a hallmark of ALS/FTD^114, 115^. We hypothesized that the nuclear chaperones DNAJC8 and DNAJC17 may play a protective role against this process. To test this possibility, we treated human (HEK293) cells with sodium arsenite for 48 hours and assessed the nuclear-to-cytoplasmic (N/C) ratio of endogenous TDP-43 (Figure 7F). As expected, sodium arsenite treatment reduced the TDP-43 N/C ratio ∼30% compared to untreated cells (Figure 7F, G). However, cells expressing DNAJC8 exhibited only an ∼11% reduction in the TDP-43 N/C ratio, and DNAJC17 completely prevented any reduction, maintaining the N/C ratio at levels comparable to the untreated control (Figure 7F, G). Thus, DNAJC8 and DNAJC17 protect against TDP-43 mislocalization under chronic stress conditions, with DNAJC17 providing the strongest defense by fully maintaining the nuclear localization of TDP-43.

### DNAJC8 and DNAJC17 directly promote TDP-43 condensates

TDP-43 undergoes phase separation in the nucleus, where it carries out essential functions^15, 116^. DNAJC8 and DNAJC17 promote nuclear retention of TDP-43 (Figure 7F, G). Thus, we hypothesized that these chaperones help maintain TDP-43 in a functional, liquid-like state. To test this idea, we purified both chaperones and performed *in vitro* reconstitution assays using TDP-43 under conditions that support condensate formation.

Given that nuclear TDP-43 condensates typically incorporate RNA^81^, we also examined the effects of nonspecific yeast total RNA and the 3.7 kb isoform of the human lncRNA NEAT1, a known paraspeckle component that interacts with TDP-43^19, 21^. To visualize condensates, we combined Alexa594-labeled and unlabeled TDP-43 at a 1:20 ratio. In buffer alone, TDP-43 formed small condensates, and at the concentrations employed neither yeast RNA nor NEAT1 (3 ng/µL) substantially altered condensate number or size (Figure 8A; S10A, B).

**Figure 8.**
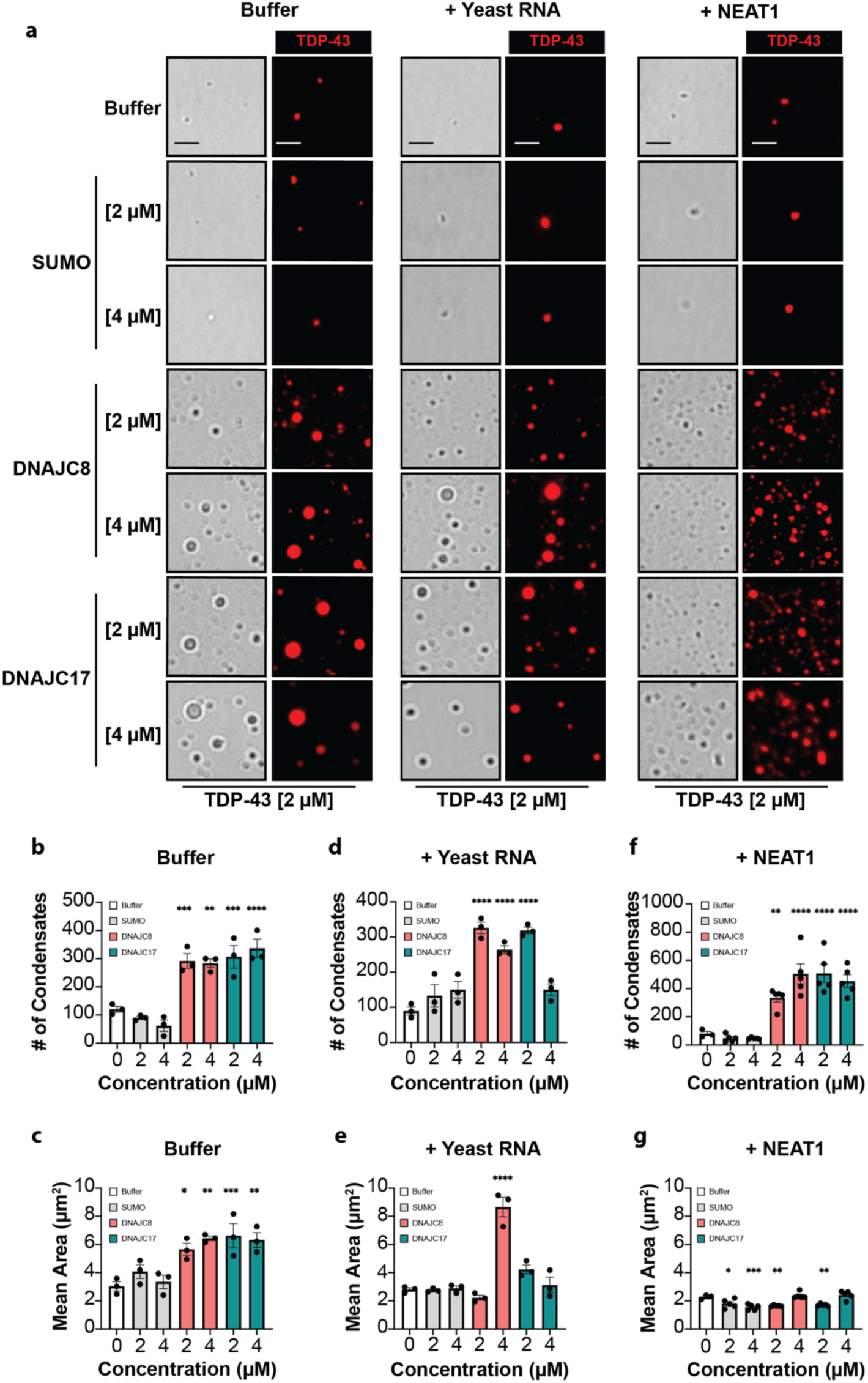
DNAJC8 and DNAJC17 promote TDP-43 condensation. **(A)** TDP-43 (95% unlabeled + 5% Alexa594-labeled) forms condensates alone and in the presence of yeast total RNA or NEAT1 lncRNA (3 ng/µL). All proteins and RNAs were diluted in assay buffer (20 mM Tris pH 7.4, 200 mM NaCl, 1 mM DTT). RNAs were used at a final concentration of 3 ng/µL. Brightfield images (left panels) and fluorescence images (right panels) show TDP-43 condensates. Scale bar, 2.5 µm. **(B)** Quantification of TDP-43 condensate number in assay bufferwith addition of SUMO, DNAJC8, or DNAJC17. Values represent mean ± SEM from three replicates. **(C)** Quantification of TDP-43 condensate area in buffer with addition of SUMO, DNAJC8, or DNAJC17. Values represent mean ± SEM from three replicates. **(D, E)** Same as (B, C), with the addition of yeast total RNA. **(F, G)** Same as (B, C), with the addition of NEAT1 lncRNA. Statistical significance was determined relative to the buffer condition by one-way ANOVA and Dunnett’s multiple comparisons test (*p < 0.05, **p < 0.01, ***p < 0.001, ****p < 0.0001).

In the absence of RNA, both DNAJC8 and DNAJC17 at 2 or 4 µM strongly enhanced TDP-43 phase separation. DNAJC8 and DNAJC17 increased both the number and size of condensates (Figure 8A–C). This effect was not observed with the control protein SUMO.

Importantly, none of the individual components (SUMO, DNAJC8, DNAJC17, yeast RNA, or NEAT1) formed condensates on their own or when combined without TDP-43 (Figure S10C). This finding confirms that the observed effects require the presence of TDP-43. It also underscores the specificity of the chaperone–RNA–TDP-43 interactions.

In the presence of yeast RNA, DNAJC8 and DNAJC17 had distinct effects on TDP-43 condensates (Figure 8A, D, E). At 2 µM, both chaperones increased condensate number compared to the buffer and SUMO, but did not strongly affect condensate size. At 4 µM, DNAJC8 increased condensate number, and triggered the formation of very large condensates, whereas DNAJC17 did not affect condensate number or size.

With NEAT1, at 2 and 4 µM both chaperones again increased TDP-43 condensate number compared to the buffer and SUMO (Figure 8A ,F , G). The chaperones showed no specific effect on the size of the NEAT1-containing TDP-43 condensates (Figure 8G). Incorporation of NEAT1 into TDP-43 condensates was confirmed using Cy5-labeled UTP (Figure S10D–F). These data indicate that the RNA component of nuclear condensates can shape the response to each chaperone.

Together, these results demonstrate that DNAJC8 and DNAJC17 promote TDP-43 phase separation in both RNA-free and RNA-rich environments. Each chaperone differentially modulates condensate properties depending on RNA context. We propose that DNAJC8 and DNAJC17 help stabilize dynamic, functional TDP-43 assemblies in the nucleus.

## Discussion

In totality, our work represents the most comprehensive functional interrogation of the human Hsp70 network against TDP-43 proteotoxicity to date. Our findings deliver a major conceptual advance in understanding how the proteostasis network controls a central driver of neurodegeneration. Indeed, we reveal a vast, previously hidden proteostatic arsenal that antagonizes TDP-43 aggregation and toxicity. A particularly striking discovery is the potent activity of DNAJC8 and DNAJC17, two poorly understood spliceosome-associated JDPs. These enigmatic chaperones act independently of Hsp70, directly modulate TDP-43 phase behavior, and define a previously unknown axis of nuclear proteostasis control. The ability of DNAJC8 and DNAJC17 to promote TDP-43 condensation and reduce aggregation in human cells enhances our understanding of how phase-separating RNA-binding proteins are regulated in health and disease. Given the urgent need for disease-modifying therapies for TDP-43 proteinopathies, our findings offer an expanded and mechanistically rich framework for therapeutic intervention—revealing previously hidden vulnerabilities in the TDP-43 aggregation pathway that are now actionable through specific proteostasis-based strategies.

To systematically uncover endogenous defenses against TDP-43 toxicity, we harnessed a highly tractable yeast model of TDP-43 proteinopathy to interrogate the entire human Hsp70 chaperone system. This comprehensive screen recapitulated known chaperone suppressors of TDP-43, including DNAJC7, which is linked to ALS/FTD^58^. Beyond these chaperones, the screen uncovered previously unrecognized suppressors with no prior connection to TDP-43 pathology. Our discovery that ∼50% of the individual components of the human Hsp70 chaperone system can buffer TDP-43 toxicity reveals a formidable proteostatic barrier that counters aberrant TDP-43 behavior. This barrier may help explain why most individuals do not develop ALS/FTD. Age-related decline in Hsp70 function^52–55^ could erode this barrier, increasing susceptibility to TDP-43 proteinopathy. Our findings illuminate the therapeutic potential in selectively boosting key components of the Hsp70 network to fortify protection against neurodegeneration. The most potent chaperones were enriched for nuclear localization, suggesting a role in safeguarding TDP-43 within the nucleus. Quantifying the aggregation burden revealed that chaperones suppress TDP-43 toxicity through multiple mechanisms. Furthermore, disease-linked and synthetic liquid variants of TDP-43 selectively evaded certain chaperones, even discriminating between highly similar isoforms such as HSPH1α and HSPH1β. Applying this approach to additional disease variants may enable personalized chaperone-based therapeutic strategies.

Genetic dissection of the yeast Hsp104 and Ssa1–4 pathways revealed that DNAJB5, DNAJB6a, DNAJB6b, and DNAJB8 rely on redundant Hsp104- and Ssa1-dependent mechanisms. DNAJB7 was an exception, potentially collaborating with distinct Hsp70s. In contrast, DNAJC8, DNAJC17, HSPA1L, HSPH1α, HSPH1β, and HSPH2 suppressed TDP-43 toxicity independently of Hsp104 and Ssa1–4, indicating noncanonical modes of action. These findings were supported by studies using mHPD JDPs and NBD-inactive HSPA1L, HSPH1α, HSPH1β, and HSPH2 mutants. Most Class B JDPs required Hsp70 activation to suppress toxicity, whereas all Class C JDPs functioned independently of Hsp70, supporting emerging evidence that Class C JDPs have diverse roles beyond Hsp70 stimulation^30^. Among cytosolic Class B JDPs, DNAJB7 retained partial suppression in the absence of Hsp70 activation. DNAJB11, an ER-resident chaperone, was the only Class B JDP to fully suppress TDP-43 toxicity without Hsp70 activation, consistent with its ability to bind clients despite J-domain inactivation^117^. The protective activity of NBD-inactive Hsp70 and Hsp110 isoforms suggests that passive chaperoning also contributes to suppression of TDP-43 toxicity. This uncoupling from canonical JDP–Hsp70–NEF ATPase cycling could present a therapeutic advantage under conditions where ATP may be limiting as in degenerating neurons^118, 119^.

We also identified minimal domains required for protective activity. In DNAJB5, DNAJB6, DNAJB7, and DNAJB8, constructs containing only the J-domain and adjacent linker retained partial suppression of TDP-43 toxicity. These fragments lack the canonical C-terminal substrate-binding domain, suggesting they likely function by stimulating Hsp70 in a general manner. The absence of toxicity may reflect autoregulatory control mediated by the linker region. These findings suggest that failure of specific JDPs to adequately activate Hsp70 may contribute to TDP-43 proteinopathy in ALS/FTD. Thus, pharmacological activation of Hsp70 to an appropriate level is anticipated to have therapeutic utility^102–104^.

Beyond Hsp70 activation, we uncovered small protective elements within the CTDs of DNAJB6a and DNAJB7, both of which contain a β-sheet motif. The CTD of DNAJB5, which adopts a β-sandwich fold, was not protective. Additionally, minimal active fragments were identified in DNAJC8 (residues 57–143) and DNAJC17 (residues 77–137). These compact, low-toxicity fragments from Class B and Class C JDPs represent promising leads for engineering chaperone therapeutics that can be readily delivered using advanced AAV technology^120–124^.

In human cells, DNAJB5, DNAJB6a, DNAJB6b, DNAJC8, DNAJC17, HSPH1α, and HSPH1β reduced insoluble TDP-43 and, in some cases, enhanced cell viability during chronic stress. DNAJC8 and DNAJC17 also promoted nuclear retention of TDP-43 under stress conditions. Disruption of phase separation within the nucleus is increasingly recognized as a pathogenic driver in ALS/FTD, where impaired phase behavior can lead to aggregation and widespread splicing defects^116^. As RNA-processing factors that localize to the nucleus, DNAJC8 and DNAJC17 are well-positioned to maintain TDP-43 solubility and functionality. Our results show that these chaperones enhance TDP-43 phase separation and prevent accumulation of insoluble forms, likely by stabilizing dynamic assemblies in the nucleus. These activities suggest early intervention points, upstream of aggregate formation, where chaperone failure may trigger disease. Understanding why these defenses collapse in ALS/FTD could reveal actionable therapeutic targets. Augmenting the concentration or activity of DNAJC8, DNAJC17, and related chaperones in vulnerable neurons may offer a powerful strategy to halt or reverse TDP-43-driven neurodegeneration. Altogether, our work defines a broad network of human chaperones that govern TDP-43 aggregation, toxicity, and phase behavior. We uncover previously unrecognized protective domains, many of which are compact and modular. These compact, modular domains represent powerful entry points for engineering chaperone-based therapeutics to reestablish nuclear TDP-43 homeostasis in ALS/FTD and related TDP-43 proteinopathies.

## Methods

### Plasmid Construction

Chaperone genes and mutants were PCR amplified from human cDNA (Gifted from Mikko Taipale) or purchased as synthetic gene fragments (IDT) with overlapping nucleotides matching the GAL1 promoter and CYC1 terminator located in a centromeric yeast plasmid (pEBGAL1). The gene fragments were cloned to pEBGAL1 by Gibson Assembly. For expression in HEK293 cells, chaperone genes were cloned to pcDNA_CMV by Gibson Assembly. For bacterial protein expression, His6-DNAJC8, His6-DNAJC17, and His6-SUMO were cloned to pEBT7 plasmid by Gibson Assembly. All plasmids were directly confirmed by sequencing.

### Yeast Transformation

The BY4741 yeast strain (ATCC, 4040002) was used in this study. To construct mNeon and TDP-43 yeast strains, the BY4741 strain was incubated at 30°C in 50mL YPD media shaking at 250rpm until OD600 reached ∼1. Cells were then transformed with pEB413GAL1_mNeon and pEB413GAL1_TDP-43 plasmids using the standard PEG and lithium acetate transformation protocol^125^. Transformed yeast clones were selected for two days at 30°C on complete synthetic medium (CSM, MP Biomedicals) lacking histidine and supplemented with 2% glucose (SD-HIS). Chaperone plasmids were transformed to these strains after their propagation in SD-HIS. Strains containing the mNeon or TDP-43 plasmid plus the chaperone plasmid were selected on double dropout synthetic media lacking histidine and uracil (JDPs), leucine (Hsp70s) or methionine (NEFs).

### Yeast Growth Assays

For yeast growth assays, clones were grown at 30°C overnight in 5 mL of the applicable double dropout media supplemented with 2% raffinose to de-repress the pGAL1 promoter. The next day, cultures were normalized to an OD_600_ of 2, serially diluted five-fold across a 96-well plate, and spotted to double dropout agar plates containing either 2% galactose or 2% glucose using a replicator pinning tool. The plates were incubated at 30°C for 65 hours before imaging.

### Yeast Microscopy

Yeast strains were grown at 30°C overnight in 5 mL of double dropout media supplemented with 2% raffinose to de-repress the pGAL1 promoter. The next day, 600 µL of the saturated raffinose cultures were used to inoculate 6 mL galactose cultures to induce production of the mNeon/TDP-43/TDP-43-YFP and chaperone. The induction cultures were grown at 30°C for 6 hours before use in microscopy or Western blotting. For microscopy, yeast were washed once with 1 M sorbitol and resuspended in 1 M sorbitol containing Hoechst (ThermoFisher, 62249, 1 µg/mL). Cells were added to the slide with a coverslip and incubated for 5 minutes before imaging with the EVOS M5000 (ThermoFisher). Images were taken at 100x magnification and processed using ImageJ.

### Yeast Western Blotting

Yeast were grown and induced the same as for microscopy. After the 6-hour induction, cells were centrifuged and pelleted at 4,000 rpm for five minutes. The pellets were resuspended in 200 µL of 0.1 M NaOH and incubated at room temperature for 10 minutes before centrifugation at 15,000 rpm for 1 minute and removal of the supernatant. Pellets were resuspended in 1X-SDS-PAGE sample buffer and boiled at 95°C for 5 minutes. Samples were separated with SDS-PAGE using a 4-20% gradient gel (Bio-Rad, 3450033) and transferred to a PVDF membrane (Millipore, IPFL00010) using a Criterion blotter wet transfer system (Bio-Rad). Membranes were blocked (Li-Cor Intercept 927-70001) for 1 hour at room temperature and then incubated with primary antibodies: anti-TDP-43 (Proteintech, 10782-2-AP, 1:5000), anti-GFP (Sigma-Aldrich, G1544, 1:1000), anti-PGK1 (Invitrogen, 459250, 1:1000), anti-FLAG (Sigma-Aldrich, F1804, 1:1000), anti-DNAJB1 (Abcam, ab223607, 1:1000), anti-DNAJB5 (Invitrogen, PA5-97670, 1:1000), anti-DNAJB6 (Proteintech, 66587-1, 1:1000), anti-DNAJB7 (Proteintech, 18540-I-AP, 1:1000), anti-DNAJB8 (Proteintech, 17071-1-AP, 1:1000), anti-DNAJB11 (ThermoFisher, 15484-1-AP, 1:1000), anti-DNAJC8 (Invitrogen, pA5-55297, 1:1000), anti-DNAJC16 (Abcam, ab122855, 1:1000), anti-DNAJC17 (Abcam, ab235350, 1:1000), anti-DNAJC18 (ThermoFisher, 25162-1-AP, 1:1000), anti-DNAJC23 (ThermoFisher, 67352-1-IG, 1:1000), anti-DNAJC25 (ThermoFisher, bs-14389R, 1:1000) for 1-3 hours at 4°C. Membranes were washed four times with PBS-T, incubated with secondary antibodies (Li-COR 680RD anti-rabbit 1:10,000; Li-COR 800CW anti-mouse 1:10,000) in blocking buffer for 1hour at room temperature, and washed again four times with PBS-T and once with PBS. Blots were imaged using an LI-COR Odyssey FC Imager.

### HEK293 Transfections

HEK293 cells (ATCC, CRL-1573) were maintained in Dulbecco’s modified Eagle’s medium (DMEM, Gibco, 11995065) enriched with 10% fetal bovine serum (FBS, HyClone, SH30910.03) and 1% penicillin-streptomycin solution (Gibco, 15140122) and incubated in a humidified incubator at 37^°^C with 5% (v/v) CO_2_. For transfection, cells were seeded into 24-well plates at a density of 2x10^5^ cells per well 24 hours prior to transfection. Lipofectamine 3000 (Invitrogen, L3000001) was used for transfection following the manufacture’s protocol using 500 ng of each co-transfected plasmid, and 0.75 µL of Lipofectamine 3000 reagent per transfection. Cells were incubated for 48 hours after the transfection before being processed for microscopy or TDP-43 solubility assays.

### Immunofluorescence Microscopy

HEK293 cells were seeded at 2x10^5^ cells per well in 24-well plates containing a poly-D-lysine-treated coverslip 24 hours prior to transfection. Cells were transfected as above. After 48 hours, the media was aspirated, and cells were washed once with PBS before fixing for 30 minutes in 4% formaldehyde in PBS. After fixing, cells were washed once with PBS and permeabilized with 0.2% Triton X-100 for 15 minutes before washing twice with PBS. V5 primary antibody (Invitrogen, SV5-Pk1) was diluted 1:100 with 2% BSA in PBS and 30 µL of primary antibody was added to each coverslip and incubated at 4°C for 1 hour. Slides were washed 15 times in PBS + 0.05% Tween-20 before being incubated with secondary antibody (Goat anti-mouse Invitrogen A32742) diluted 1:1000 with 2% BSA in PBS for 30 minutes. Slides were washed 15 times in PBS + 0.05% Tween-20 and assembled with Mounting Medium containing DAPI (VECTASHIELD Antifade, Vector Laboratories, H-1200-10) and sealed before imaging. Images were taken at 100x magnification using the EVOS M5000 Imager (ThermoFisher) and processed using ImageJ.

### TDP-43 Solubility Assay

48 hours after transfection of 2x10^5^ cells, the cells were washed once with PBS, then resuspended in 200µL RIPA lysis buffer (150 mM NaCl, 1% Triton X-100, 1% sodium deoxycholate, 0.1% SDS, 25 mM Tris–HCl pH 7.6). Cells were then sonicated and centrifuged for 30 min at 15,000 rpm. The supernatant was removed, and the pellet was washed once with 200 µL RIPA and centrifuged again for 30 min at 15,000 rpm. The pellet was resuspended in urea buffer (8 M urea, 2 M thiourea, 4% CHAPS). 10 µL of RIPA and UREA fractions were mixed with 5 µL of 3X SDS-PAGE sample buffer. Only RIPA samples were then boiled. All samples were separated by SDS-PAGE (4–20% gradient, Bio-Rad 3450033) and transferred to a PVDF membrane (Millipore IPFL00010) using a Criterion blotter wet transfer system (Bio-Rad). Membranes were blocked (Li-Cor Intercept 927-70001) for 1 hour at room temperature and then incubated with primary antibodies: rabbit anti-TDP-43 polyclonal (Proteintech 10782-2-AP); rabbit anti-GFP polyclonal (Sigma-Aldrich G1544); mouse anti-V5 (Invitrogen SV5-Pk1) for 1-3 hours at 4°C. Membranes were washed four times with PBS-T, incubated with secondary antibodies in blocking buffer for 1hour at room temperature, and washed again four times with PBS-T and once with PBS. Blots were imaged using an LI-COR Odyssey FC Imager.

### HEK293 Cell Viability Assay

2x10^4^ HEK293 cells were seeded in 96-well plates and transfected as above using 100 ng of each plasmid. At 24 hours post-transfection, the cells were treated with 5 µM sodium arsenite (Sigma-Aldrich, S7400) and further incubated for 48 hours. To measure viability the cells were assayed with CellTiter-Glo 2.0 Assay (Promega, G9241) according to the manufacturer’s protocol.

### HEK293 Nuclear/Cytoplasmic TDP-43 Assay

HEK293 cells were seeded in 8-well chamber slides (Lab-Tek, C7182) coated with poly-L-ornithine (Sigma-Aldrich, P4957) and allowed to attach for 24 hours. Wells were transfected with 250ng of plasmid DNA using Lipofectamine 3000 (Invitrogen, L3000001). 24 hours post-transfection, wells were treated with 5 µM sodium arsenite (Sigma-Aldrich, S7400) for 48 hours. Wells were washed with PBS three times, fixed with methanol, and permeabilized with 0.1% Triton-X 100 (ThermoScientific, A16046-AE) for 10 min. Wells were washed again with PBS three times and blocked for 1 hour at 4°C in 10% goat serum. Wells were incubated with primary antibodies (G3BP1 Mouse Monoclonal (Proteintech, 66486-1-Ig, 1:500), TDP-43 Rabbit Polyclonal (Proteintech, 10782-2-AP, 1:400), V5 Tag Mouse Monoclonal (Invitrogen, SV5-Pk1, 1:250)) overnight at 4°C. Wells were washed with PBS three times and incubated with 4 µg/mL secondary antibody in 2% goat serum for 1 hour at RT (GαR AlexaFluor 488 (Invitrogen, A-11008), GαM AlexaFluor 568 (Invitrogen, A-11004)). Wells were washed with PBS three times with Hoechst 33342 (1:5000, Invitrogen, H3570) added in the second wash. Slides were mounted in fluorogel (Electron Microscopy Sciences, 17985-10), coverslipped, and imaged with confocal microscopy (Leica SP8).

### Purification of DNAJC8, DNAJC17, and SUMO

DNAJC8, DNAJC17, and SUMO were purified as N-terminal 6xHis-tagged proteins. pEBT7 plasmids encoding these proteins were transformed to BL21Star (DE3) One Shot competent *E. coli* cells using standard heat shock transformation. After recovery the bacteria were plated to LB + 100 µg/mL ampicillin (LB-Amp) plates. After incubation overnight at 37°C, colonies were harvested from the plate and used to begin a 200 mL LB-Amp starter culture. After 3-4 hours, 30mL of starter culture was added to 6 large 1 L LB-Amp cultures and these were shaken at 250 rpm in 37°C until OD_600_ reached ∼0.8-1. The cultures were cooled at 4°C then induced with 1 mM IPTG for 16 hours at 16°C. Cells were centrifuged at 4,000 rpm for 20 minutes and the pellets were each resuspended in 20 mL of lysis buffer (20 mM Tris pH 7.4, 1 M NaCl, 10% Glycerol, 1 mM DTT, 5 µM pepstatin A, 100 µM PMSF, 10 µg/mL RNAse, 10 µg/mL DNAse, 10 mM imidazole, and 1 tablet/50 mL cOmplete EDTA-free Protease Inhibitor Cocktail (Roche, 5056489001) and sonicated until lysed. The lysate was centrifuged at 20,000 rpm for 45 minutes and the cleared lysate was incubated with rotation in 12 mL of Ni-NTA affinity resin (QIAGEN, 30250) that was pre-equilibrated with lysis buffer, for 1 hour at 4°C. Beads were then washed three times with wash buffer (20 mM Tris pH 7.4, 1 M NaCl, 10% Glycerol, 1 mM DTT, 30 mM imidazole) and incubated 30 minutes with elution buffer (20 mM Tris pH 7.4, 1 M NaCl, 10% Glycerol, 1 mM DTT, 300 mM imidazole). The eluted protein was concentrated to 5 mL using an Amicon Ultra-15 Centrifugal Filter (Millipore, UFC9030 for DNAJC8/DNAJC17 and UFC9010 for SUMO ) by centrifugation at 4,000 rpm at 4°C, then loaded for size exclusion chromatography on a HiLoad 16/600 Superdex 75 pg column (Cytiva, 28989333) pre-equilibrated in size exclusion buffer (20 mM Tris pH 7.4, 1 M NaCl, 10% Glycerol, 1 mM DTT) and eluted fractions containing pure protein were analyzed via SDS-PAGE to identify the protein based on size. Pure fractions containing the protein of interest were pooled, then concentrated using an Amicon Ultra-15 Centrifugal Filter (Millipore, UFC9030 for DNAJC8/DNAJC17 and UFC9010 for SUMO) until a concentration of >200 µM was achieved. Aliquots of the protein were flash-frozen in liquid nitrogen and stored at -80°C until use.

### Purification of TDP-43-MBP-his6 for phase separation assays

TDP-43-MBP-his6 expression plasmid was transformed to BL21Star (DE3) One Shot competent *E. coli* cells using standard heat shock transformation. After recovery the bacteria were plated to LB + 50 µg/mL kanamycin (LB-Kan) plates. After incubation overnight at 37°C, colonies were harvested and used to begin six 1 L LB-Kan cultures supplemented with 0.2% dextrose shaken at 250 rpm at 37°C until OD_600_ reached ∼0.5-0.9. The cultures were cooled at 4°C then induced with 1 mM IPTG for 16 hours at 16°C. Cells were centrifuged at 4,000 rpm for 20 minutes and the pellets were each resuspended in 20 mL of lysis buffer (20 mM Tris pH 8, 1 M NaCl, 10% Glycerol, 1 mM DTT, 10 mM imidazole, and 1 tablet/50 mL cOmplete EDTA-free Protease Inhibitor Cocktail (Roche, 5056489001) and sonicated until lysed. The lysate was then centrifuged at 20,000 rpm at 4°C for 45 min and the cleared lysate was filtered and loaded to a HisTrap column (5 mL, Cytiva, 17524801) pre-equilibrated in lysis buffer. The column was then washed with 20 mL wash buffer (20 mM Tris pH 8, 1 M NaCl, 10% Glycerol, 1 mM DTT, 30 mM imidazole) and eluted with 0-80% gradient elution using elution buffer (20mM Tris pH 8, 1M NaCl, 10% Glycerol, 1mM DTT, 500 mM imidazole). The eluted sample was then pooled and spin concentrated to <13 mL using an Amicon Ultra-15 Centrifugal Filter MWCO 50 kDa (Millipore, UFC9050) before loading to a size exclusion column 26/600 Superdex 200 pg column (Cytiva, 28989335) pre-equilibrated in SEC buffer (20 mM Tris pH 8, 300 mM NaCl). The second peak as evaluated by absorbance at 280nm was collected and concentrated using an Amicon Ultra-15 Centrifugal Filter MWCO 50 kDa (Millipore, UFC9050) until a concentration of >200 µM was achieved. Aliquots were flash-frozen in liquid nitrogen and stored at -80°C until use.

### Purification of TEV protease

His6-TEV plasmid was transformed to BL21Star (DE3) One Shot competent *E. coli* cells with standard heat shock and plated to LB + 100 µg/mL ampicillin (LB-Amp) plates. After incubation overnight at 37°C, colonies were harvested and used for a 50mL LB-Amp starter culture. After 2 hr, the starter culture was diluted 1:100 into a 1L LB-Amp culture of LB and shaken at 250 rpm and 37°C until the OD_600_ reached ∼0.7, then cooled at 4°C and induced with 1 mM IPTG (MilliporeSigma, 420322), and grown shaking at 250 rpm for 16 hours at 16°C. After induction, the culture was harvested by centrifugation at 4,000 rpm at 4°C for 20 min. The pelleted cells were resuspended in 30 mL Lysis Buffer (25 mM Tris-HCl pH 8.0, 500 mM NaCl, 1 mM DTT, and 1 tablet/50 mL of cOmplete, EDTA-free Protease Inhibitor Cocktail (MilliporeSigma, 5056489001)). The lysate was then centrifuged at 20,000 rpm at 4°C for 45 min and the cleared lysate was incubated with rotation in 2mL of Ni-NTA affinity beads (QIAGEN, 30250) that were pre-equilibrated with lysis buffer, for 1 hour at 4°C then centrifuged at 2,000 rpm at 4°C for 5 min. The Ni-NTA resin was then washed with 25 column volumes (CV) of Wash Buffer (25 mM Tris-HCl pH 8.0, 500 mM NaCl, 1 mM DTT, 25 mM imidazole), with centrifugations performed at 2,000 rpm at 4°C for 5 min. Protein was eluted with 5 CV of Elution Buffer (25 mM Tris-HCl pH 8.0, 500 mM NaCl, 1 mM DTT, 300 mM imidazole). Eluted protein was concentrated to 5 mL using an Amicon Ultra-15 Centrifugal Filter Unit, MWCO 10 kDa (Millipore, UFC9010), by centrifugation at 4,000 rpm at 4°C and then was loaded for size exclusion chromatography on a 16/600 Superdex 200 pg column (Cytiva, 28989335) pre-equilibrated in size exclusion buffer (25mM Tris pH 7, 300mM NaCl, 10% Glycerol, 1mM DTT) and eluted fractions containing pure protein were analyzed via SDS-PAGE. Pure fractions were pooled, then concentrated using an Amicon Ultra-15 Centrifugal Filter (Millipore, UFC9010) until a concentration of >10 mg/mL was achieved. Aliquots were flash-frozen in liquid nitrogen and stored at -80°C until use.

### In vitro transcription of NEAT1 lncRNA

The 3.7 kb NEAT1_1 lncRNA isoform was transcribed with MEGAscript in vitro transcription kit (ThermoFisher, AM1334) using pCRII_NEAT-short_IVT plasmid linearized with restriction digest by BamHI-HF (New England Biolabs, R0136). To label the RNA with Cy5 for visualization, 1 µL of Cy5-labeled UTP was added to the reaction according to manufacturer specifications.

### Alexa594 dye labeling of TDP-43-MBP-his6

TDP-43-MBP-his6 was equilibrated in 150 mM NaCl, 20 mM HEPES pH 7.4, 1 mM DTT using a Micro Bio-Spin P-6 Gel column (Bio-RAD, 7326200) before labeling. Alexa594 dye (ThermoFisher, A20004) was dissolved in DMSO at a concentration of 10 mg/mL, mixed 1:50 with TDP-43-MBP-his6, and incubated at room temperature in the dark for 1 hour. After incubation, the free dye was removed by filtering through two Micro Bio-Spin P-6 Gel columns (Bio-RAD, 7326200). Aliquots of the labeled protein were flash-frozen in liquid nitrogen and stored at -80°C until use.

### TDP-43 phase separation assay

TDP-43-MBP-his6, DNAJC8, DNAJC17, SUMO, RNA, and TEV protease were thawed on ice for 10 minutes before use. TDP-43 was centrifuged at 15,000 rpm for 10 minutes. All proteins and RNAs were diluted in assay buffer (20 mM Tris pH 7.4, 200 mM NaCl, 1 mM DTT). TDP-43-MBP-his6 was prepared with yeast total RNA (Roche, 10109223001) or NEAT1 to achieve 2 µM TDP-43 final concentration and 3 ng/µL final RNA concentration. Equal volumes of diluted TDP-43-MBP-his6 (±RNA) and chaperones (at variable concentrations) were combined and incubated at room temperature for 10 minutes before an equal volume of TEV protease (30 µg/mL final concentration) was added to start the TDP-43 phase separation reaction. Reactions were placed sealed on a microscope slide and incubated at room temperature for 1 hour before imaging at 100x magnification with the EVOS M5000 Imager (ThermoFisher). Condensates were quantified using CellProfiler software.

### Quantification and Statistical Analysis

Yeast growth was quantified using CellProfiler software. The yeast images were converted to grayscale and the yeast spots were identified in a grid using the adaptive Otsu three class threshold method with an adaptive window size of 150 pixels and a threshold correction factor of 1.3. Yeast growth was measured using the MeasureObjectIntensity module.

Chaperone subcellular compartment associations (Table S1) were compiled from UniProt (www.uniprot.org) and Piette *et al*.^30^. For each growth phenotype (e.g., toxic with mNeon), the percentage of hit chaperones localized to each compartment was calculated and compared to the overall library to determine percent enrichment. Statistical significance for enrichment within individual compartments was assessed using a chi-square test on a 2×2 contingency table with GraphPad Prism software, comparing the number of hits and non-hits associated with each compartment.

Quantification of yeast TDP-43-YFP foci number and size was performed with CellProfiler using the IdentifyPrimaryObjects module with a minimum size of 1 pixel and maximum size of 50 pixels. Objects outside of this range were discarded. The global Otsu two class threshold method was used to identify the foci and the MeasureObjectSizeShape module was used to determine the foci area. Quantification of nuclear and cytoplasmic TDP-43 in HEK293 cells was performed with CellProfiler. DAPI was used to define the nucleus with the IdentifyPrimaryObjects module. The diameter range was set to 30-150 pixels. Objects outside of this range were discarded. The global minimum cross-entropy threshold method was used with a smoothing scale of 1.3488. Diffuse G3BP1 signal defined the cell boundary using the propagation setting in the IdentifySecondaryObjects module and global minimum cross-entropy threshold method with a smoothing scale of 1.3488. The IdentifyTertiaryObjects method was used to define the cytoplasm as the subtraction of the nucleus from the cell boundary area. TDP-43 signal was measured in both the nucleus and cytoplasm using the MeasureObjectIntensity module.

TDP-43 solubility in RIPA and urea fractions was quantified by measuring Western blot band intensities using the rectangle selection tool in Image Studio Lite software (LI-COR Biosciences).

Quantity and size of purified TDP-43 condensates was measured using CellProfiler software using the IdentifyPrimaryObjects module with a minimum size of 1 pixel and maximum size of 40 pixels. The adaptive Otsu two class threshold method with an adaptive window of 50 pixels was used to identify the condensates and the MeasureObjectSizeShape module was used to determine condensate area.

All statistical analyses were performed with GraphPad Prism software. Details for statistical tests in each experiment are outlined in the figure legends.

## Acknowledgements

We thank Mikko Taipale for generous provision of plasmids. We thank JiaBei Lin, Linamarie Miller, Zarin Tabassum, Zoe Chen, Clarice Xu, and Sabrina Lin for feedback on the manuscript. E.M.B. was supported by a Milton Safenowitz Post-Doctoral Fellowship from ALS Association and NIH grants F32NS108598 and K99AG075242. M.L. was supported by a Milton Safenowitz Post-Doctoral Fellowship from ALS Association, a Mildred Cohn Distinguished Postdoctoral Award, and an Alzheimer’s Association Research Fellowship. K.E.C. was supported by NIH grants T32GM132039 and F31NS129101. D.A.A. and K.R.W. were supported by NIH grant K08NS114106. B.L.D. was supported by the Children’s Hospital of Philadelphia Research Institute. J.S. was supported by The Packard Center for ALS Research at Johns Hopkins, Hop on a Cure Foundation, Target ALS, The Association for Frontotemporal Degeneration, ALS Association, G. Harold and Leila Y. Mathers Foundation, Alzheimer’s Association Zenith Research Fellows Award, Kissick Family Foundation and the Milken Institute Science Philanthropy Accelerator for Research and Collaboration, Department of Defense grants W81XWH-17-1-0237 and W81XWH-20-1-0242, and NIH grants R21AG065854 and R01GM099836.

## Author contributions

Conceptualization, E.M.B., M.L., D.A.A., & J.S.; methodology, E.M.B., M.L., K.R.W., Y.C., S.B., K.E.C., D.A.A., & J.S.; validation, E.M.B., M.L., K.R.W., Y.C., S.B., P.M.C., B.L., & K.V.; formal analysis, E.M.B., M.L., K.R.W., D.A.A., & J.S.; investigation, E.M.B., M.L., K.R.W., Y.C., S.B., P.M.C., B.L., & K.V.; software, E.M.B.; resources, E.M.B., M.L., K.R.W., K.E.C., D.A.A., B.L.D., & J.S.; data curation, E.M.B., K.R.W., D.A.A., B.L.D., & J.S.; writing– original draft, E.M.B., & J.S.; writing – review & editing, E.M.B., M.L., K.R.W., Y.C., S.B., P.M.C., B.L., K.V., D.A.A., B.L.D., & J.S.; visualization, E.M.B., K.R.W., D.A.A., & J.S.; supervision, E.M.B., D.A.A., B.L.D., & J.S.; project administration, E.M.B., D.A.A., B.L.D., & J.S.; funding acquisition, E.M.B., M.L., K.E.C., D.A.A., B.L.D., & J.S.

---

**Table S1. Ranked summary of Hsp70 network components screened for suppression of TDP-43 toxicity in yeast**.Proteins are listed in order of decreasing growth rescue (TDP-43 Mean Growth/Vector). Chaperones with a statistically significant difference from vector control (p < 0.05) determined by one-way ANOVA and Dunnett’s multiple comparisons test. The table includes each protein’s UniProt-reported subcellular localization, and its compartment association based on experimental data^30^.

**Supplementary Figure 1.**
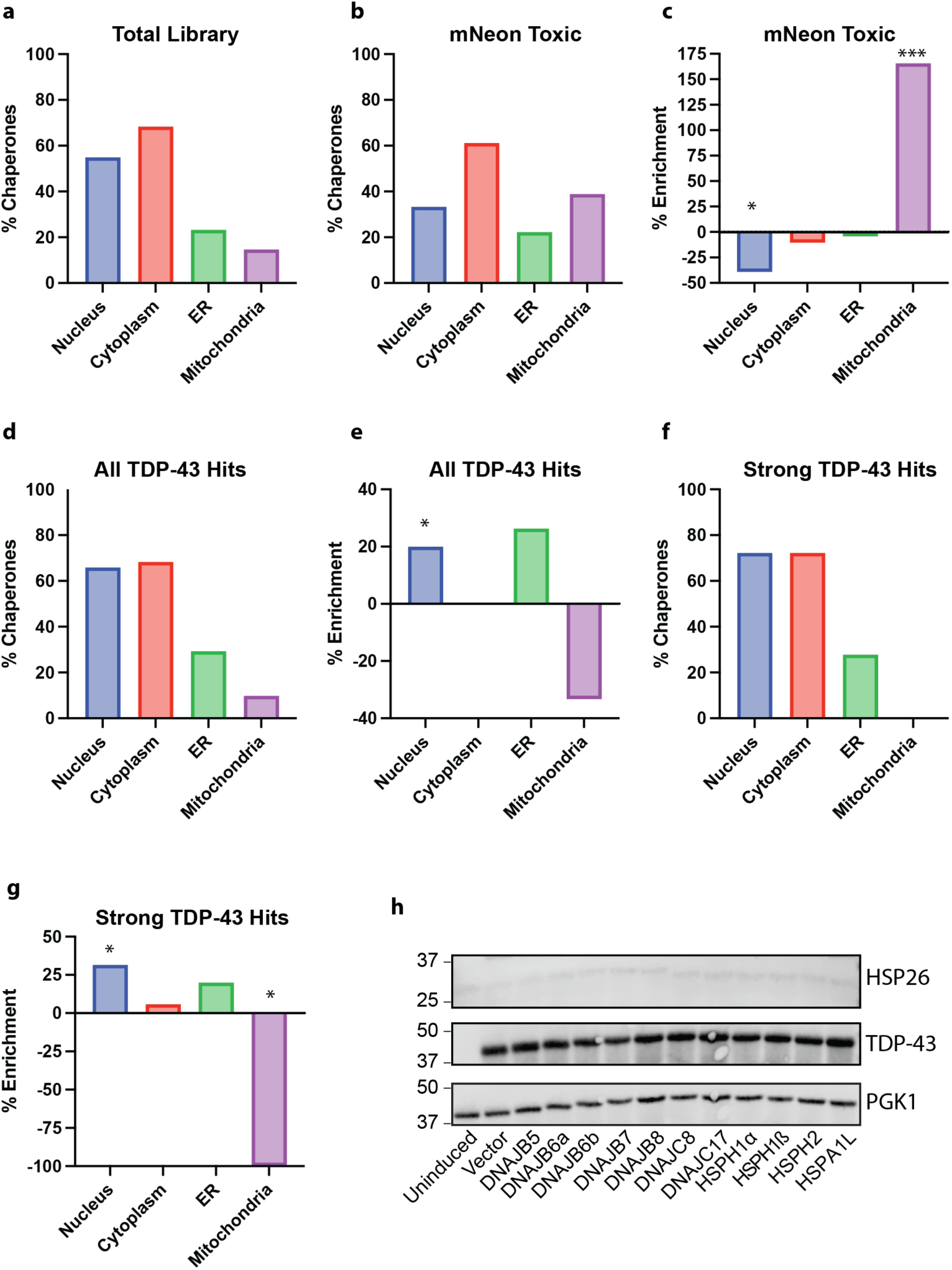
Components of the human Hsp70 network that mitigate TDP-43 toxicity localize to the nucleus, ER, and cytoplasm and do not induce a general stress response. (**A**) Distribution of subcellular localization among all human Hsp70 network components included in the screen, categorized as nuclear, cytoplasmic, ER, or mitochondrial (%). The total exceeds 100% because many chaperones are associated with multiple subcellular compartments (see Table S1). (**B**) Distribution of subcellular localization for human Hsp70 network components that reduced growth by >10% in the mNeon-expressing control strain (%). (**C**) Enrichment or depletion (%) of human Hsp70 network components from each subcellular compartments that impaired growth in the mNeon control strain relative to the total library. (**D**) Distribution of subcellular localization for the 41 human Hsp70 network components that enhanced growth in the TDP-43-expressing strain (%). (**E**) Enrichment or depletion (%) of the 41 human Hsp70 network components from each subcellular compartment that enhanced growth in the TDP-43 strain relative to the total library. (**F**) Distribution of subcellular localization for human Hsp70 network components that strongly enhanced growth (>50%) in the TDP-43-expressing strain (%). (**G**) Enrichment or depletion (%) of human Hsp70 network components from each subcellular compartment that strongly enhanced growth (>50%) in the TDP-43 strain relative to the total library. Statistical significance for enrichment within individual compartments was assessed by chi-square test comparing the number of hits and non-hits associated with each compartment (*p < 0.05, ***p < 0.001). (**H**) Western blot images for PGK1 (loading control), TDP-43, and HSP26 expression in strains harboring TDP-43 and the indicated chaperones.

**Supplementary Figure 2.**
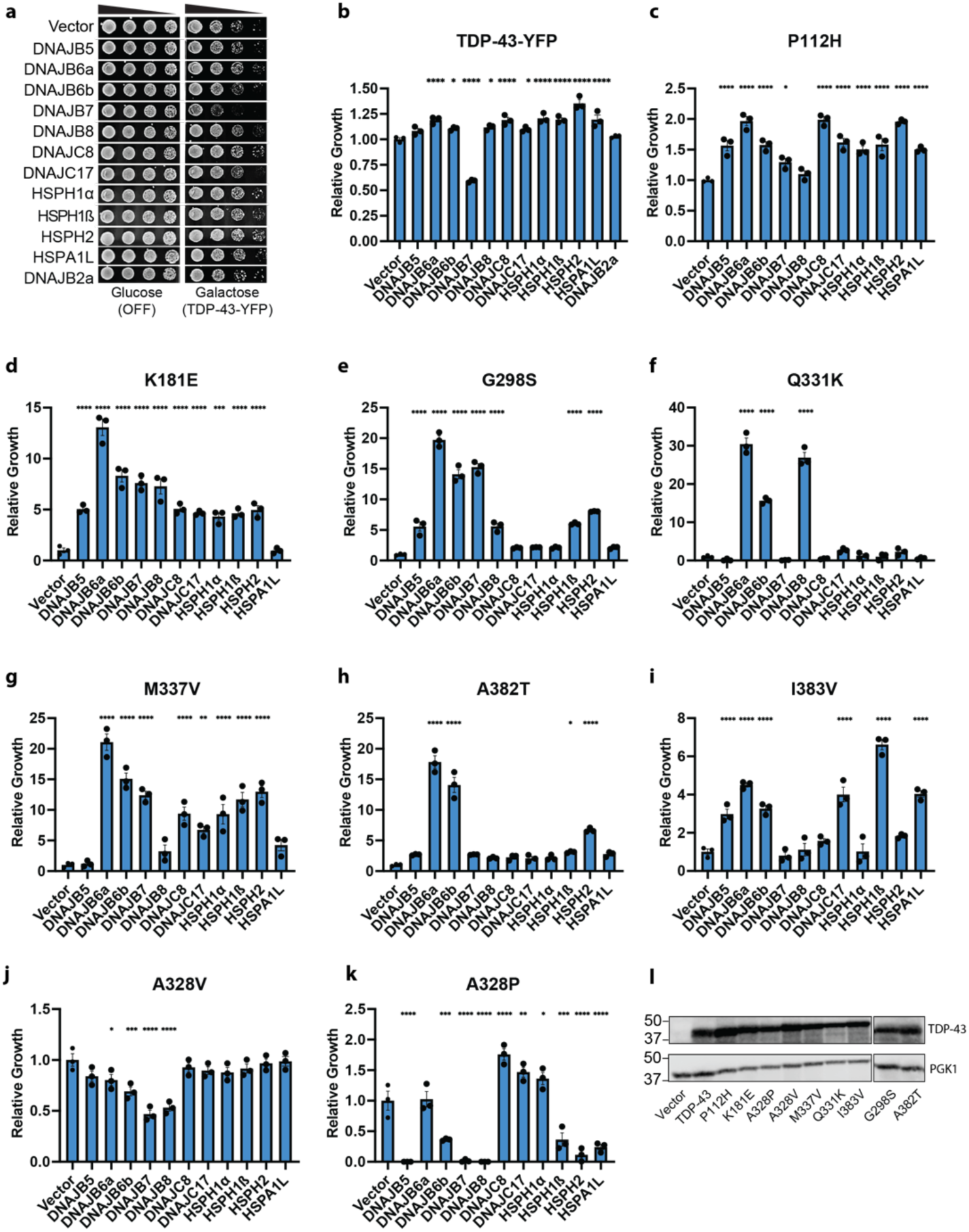
Human chaperones buffer the toxicity of diverse synthetic and disease-linked TDP-43 variants. (**A**) Yeast strains harboring galactose-inducible TDP-43-YFP were transformed with plasmids encoding galactose-inducible human chaperones. On glucose media, there is no expression of TDP-43-YFP or chaperones. Cultures were normalized to equivalent density (OD_600_ = 2), serially diluted 5-, 25-, and 125-fold, and spotted onto glucose and galactose agar plates. Images show representative yeast growth. (**B**) Quantification of relative growth normalized to the vector control. Values represent mean ± SEM from three independent replicates. (**C-K**) Quantification of relative growth normalized to the vector control for each indicated TDP-43 variant against the panel of human chaperones. Values represent mean ± SEM from three independent replicates. Statistical significance was calculated relative to the vector control using one-way ANOVA and Dunnett’s multiple comparisons test (*p < 0.05, **p < 0.01, ***p < 0.001, ****p < 0.0001). (**L**) Western blot images for individual TDP-43 variants confirming their expression. PGK1 is used as a loading control.

**Supplementary Figure 3.**
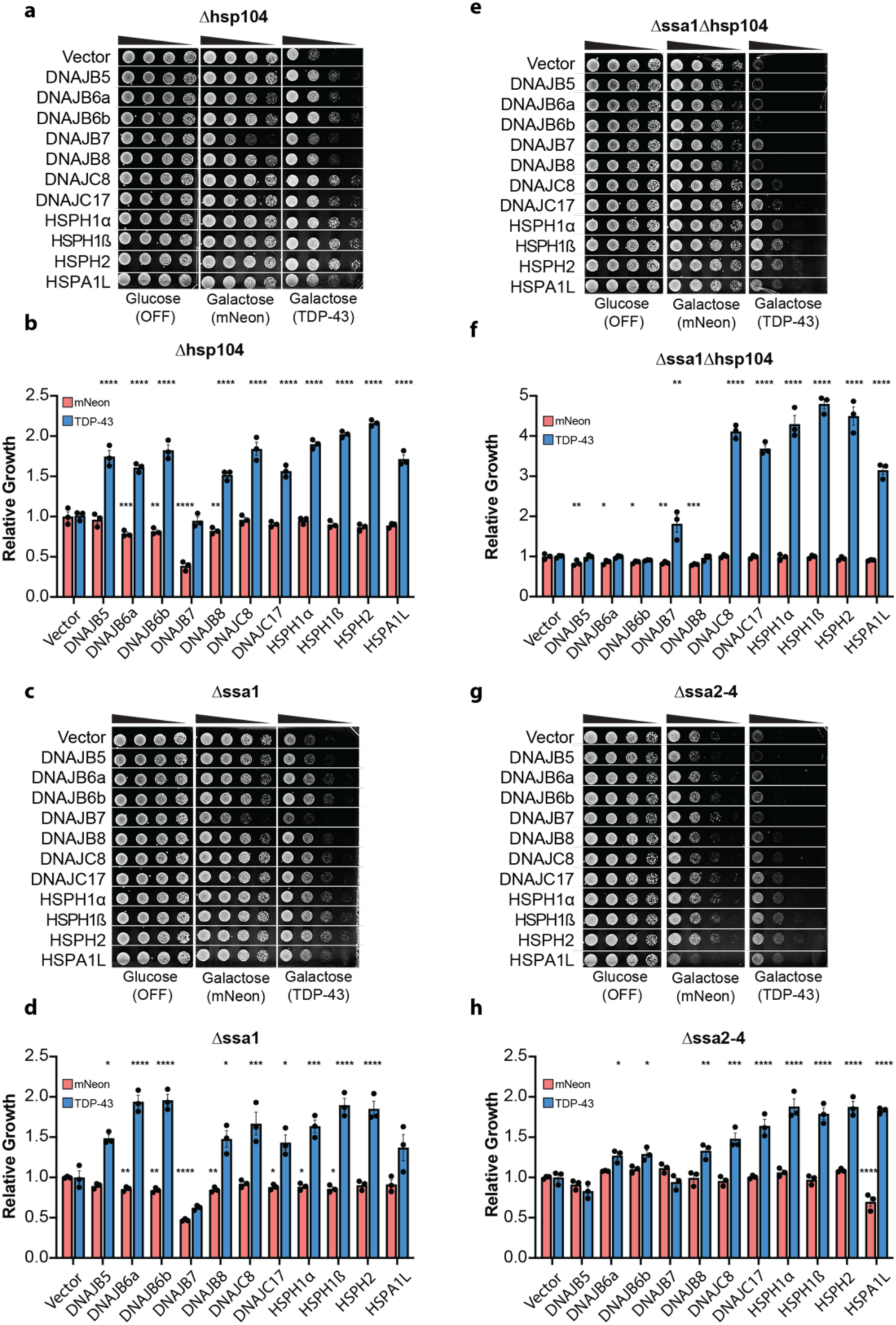
DNAJC8, DNAJC17, HSPA1L, HSPH1α, HSPH1β, and HSPH2 suppress TDP-43 toxicity independently of Hsp104 and Ssa1-4. (**A**) Δhsp104 yeast strains harboring galactose-inducible mNeon or TDP-43 were transformed with plasmids encoding galactose-inducible human chaperones. On glucose media, there is no expression of mNeon, TDP-43, or chaperones. Cultures were normalized to equivalent density (OD_600_ = 2), serially diluted 5-, 25-, and 125-fold, and spotted onto glucose and galactose agar plates. Images show representative yeast growth. (**B**) Quantification of relative growth normalized to the vector control. Values represent mean ± SEM from three independent replicates. (**C-H**) Same as (A, B) for Δssa1 (C, D), Δ*hsp104*Δ*ssa1* (E, F), Δ*ssa2-4* (G, H). Statistical significance was calculated relative to the vector control using one-way ANOVA and Dunnett’s multiple comparisons test (*p < 0.05, **p < 0.01, ***p < 0.001, ****p < 0.0001).

**Supplementary Figure 4.**
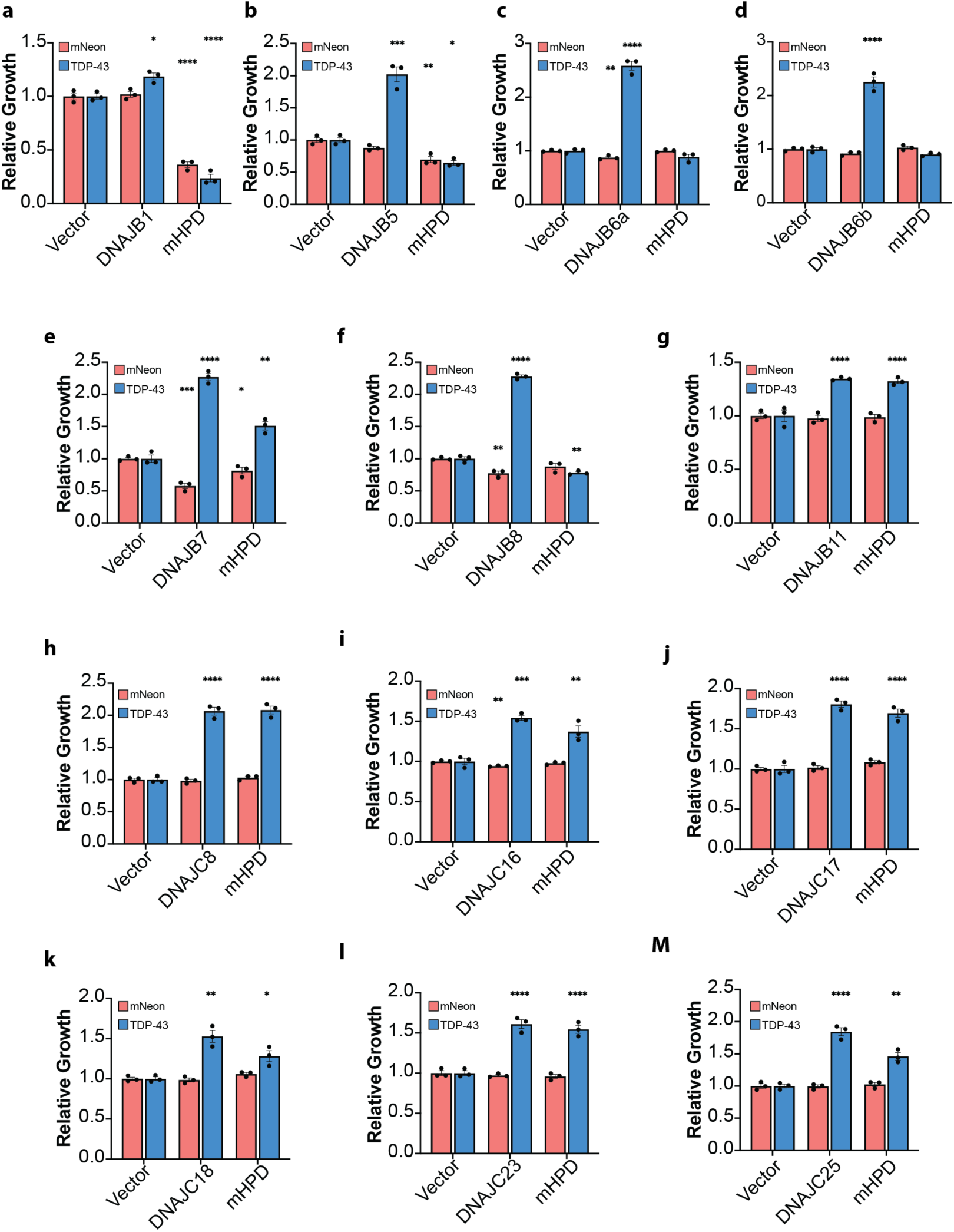
HPD motif mutants reveal Hsp70-dependent and Hsp70-independent JDPs. (**A-M**) Quantification of relative yeast growth for indicated JDP and corresponding mHPD against mNeon or TDP-43. Values represent mean ± SEM of three replicates. Statistical significance was calculated relative to the vector control using one-way ANOVA and Dunnett’s multiple comparisons test (*p < 0.05, **p < 0.01, ***p < 0.001, ****p < 0.0001).

**Supplementary Figure 5.**
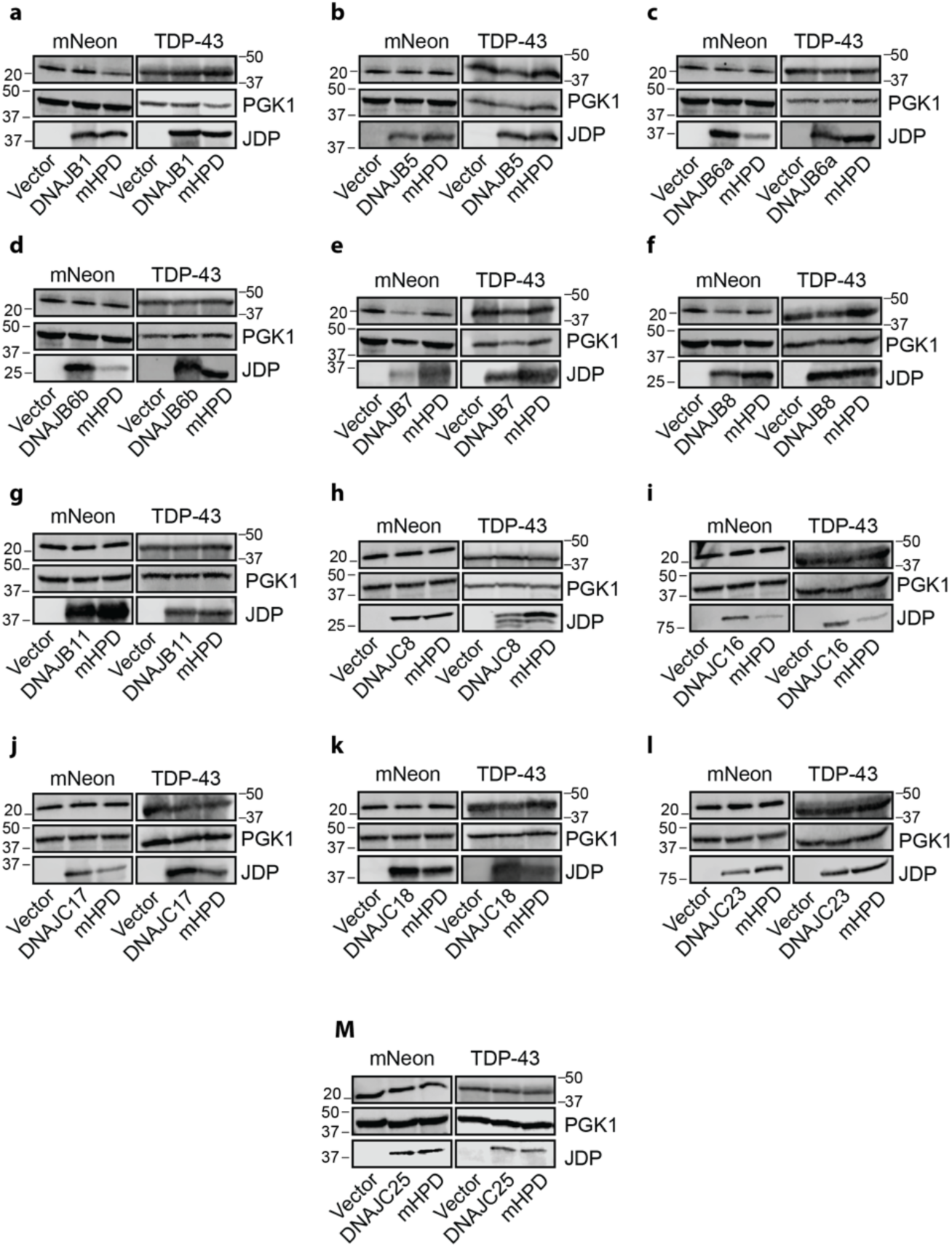
Western blots for JDPs and mHPDs. (**A-M**) TDP-43 and mNeon Western blots for the indicated JDP and corresponding mHPD. PGK1 is used as a loading control. Molecular weight markers are indicated.

**Supplementary Figure 6.**
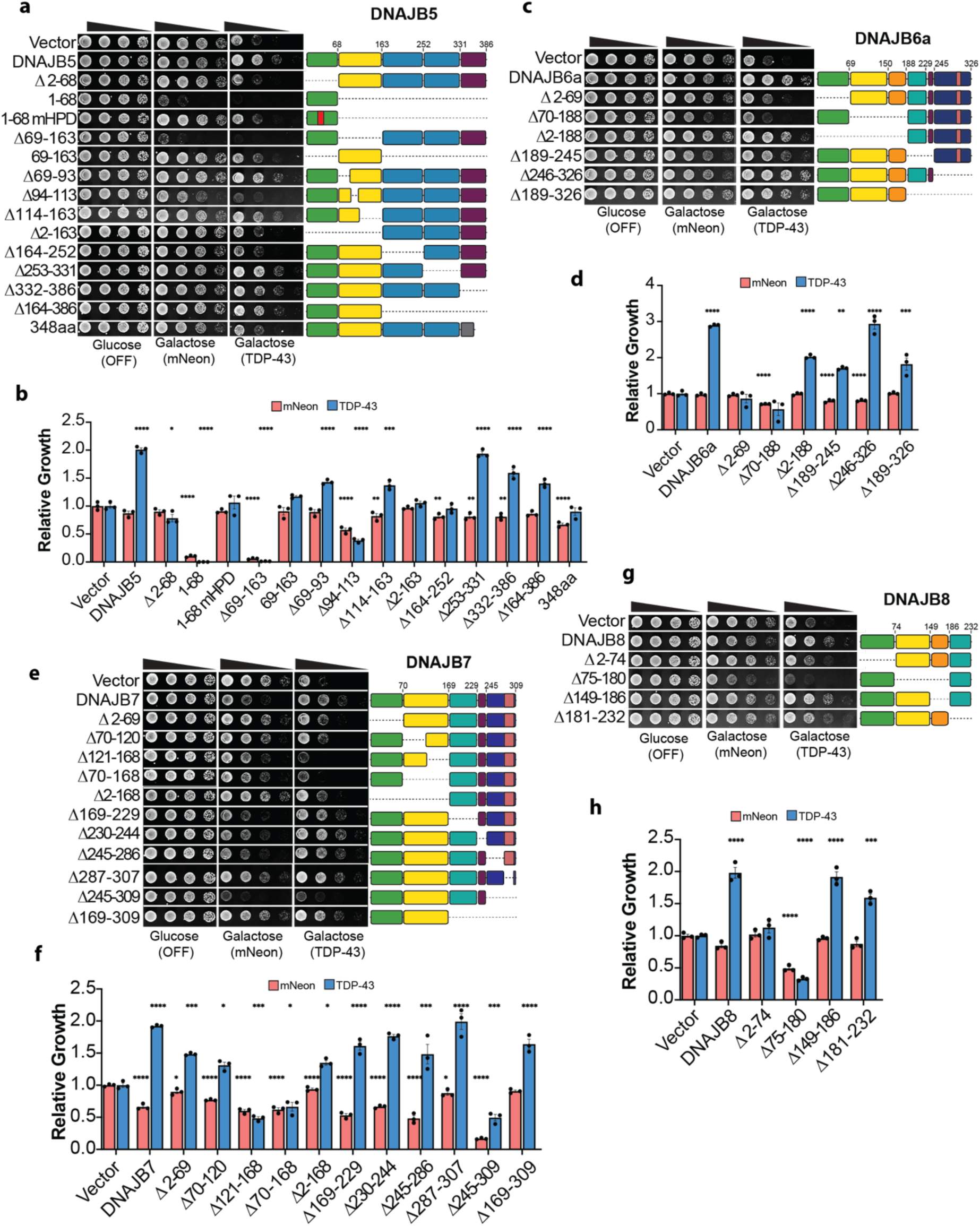
Domain deletion analysis for Class B JDPs. (**A**) Yeast strains harboring mNeon or TDP-43 were transformed with plasmids encoding DNAJB5 domain deletion constructs. Strains were grown on glucose (no mNeon, TDP-43, or chaperone expression) or galactose (induced expression) plates. Cultures were normalized to equivalent density (OD_600_ = 2), serially diluted 5-, 25-, and 125-fold, and spotted onto glucose and galactose plates. Representative yeast growth assay images for DNAJB5 mutants. (**B**) Quantification of relative yeast growth for DNAJB5 mutants. (**C-H**) Same as A,B for DNAJB6a (**C,D**), DNAJB7 (**E,F**), and DNAJB8 (**G,H**). Values are mean ± SEM from three independent replicates. Statistical significance was determined relative to the vector control by one-way ANOVA and Dunnett’s multiple comparisons test (*p < 0.05, **p < 0.01, ***p < 0.001, ****p < 0.0001).

**Supplementary Figure 7.**
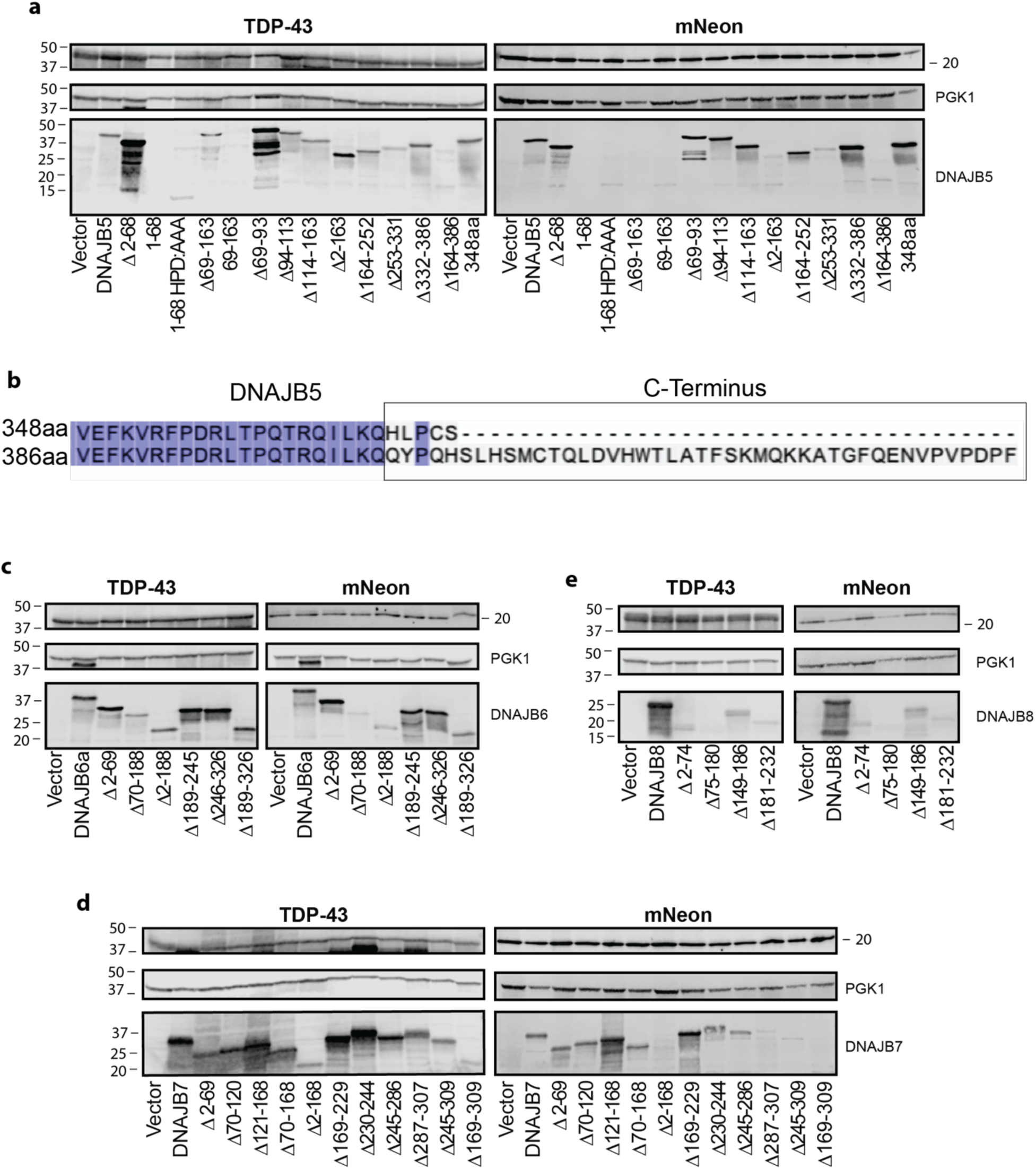
Western blots and splice isoform alignment for Class B JDP deletion mutants. (**A**) TDP-43 and mNeon Western blots for DNAJB5 mutants. PGK1 is used as a loading control. (**B**) Alignment of C-termini for two DNAJB5 splice variants. Colored amino acids are identical between both isoforms. (**C-E**) TDP-43 and mNeon Western blots for DNAJB6a (C), DNAJB7 (D), and DNAJB8 (E) mutants.

**Supplementary Figure 8.**
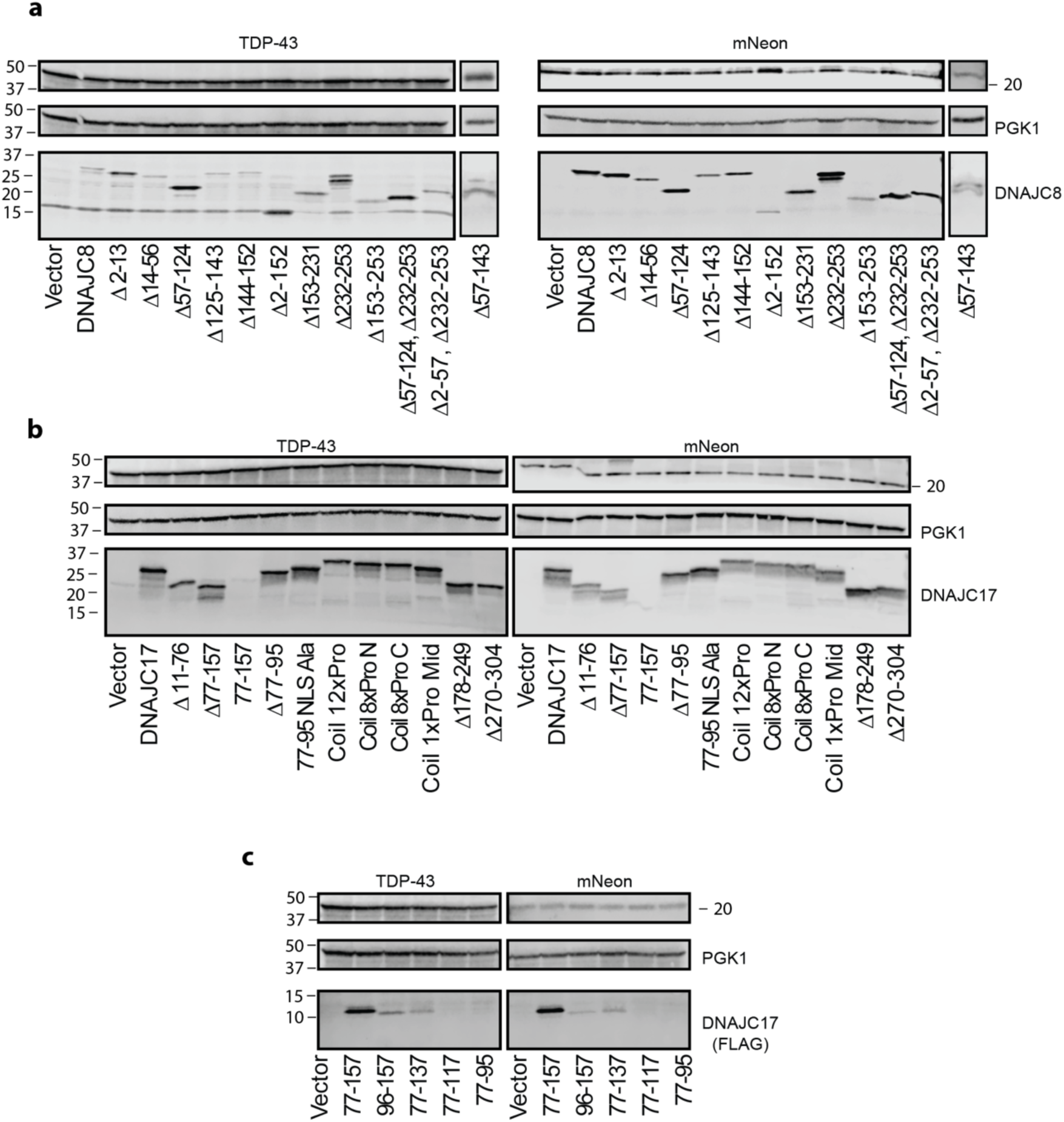
Western blots for DNAJC8 and DNAJC17 variants. (**A,B**) TDP-43 and mNeon Western blots for DNAJC8 (A) and DNAJC17 (B) mutants. (**C**) Western blot detection of FLAG tagged DNAJC17 coiled coil variants. PGK1 is used as a loading control.

**Supplementary Figure 9.**
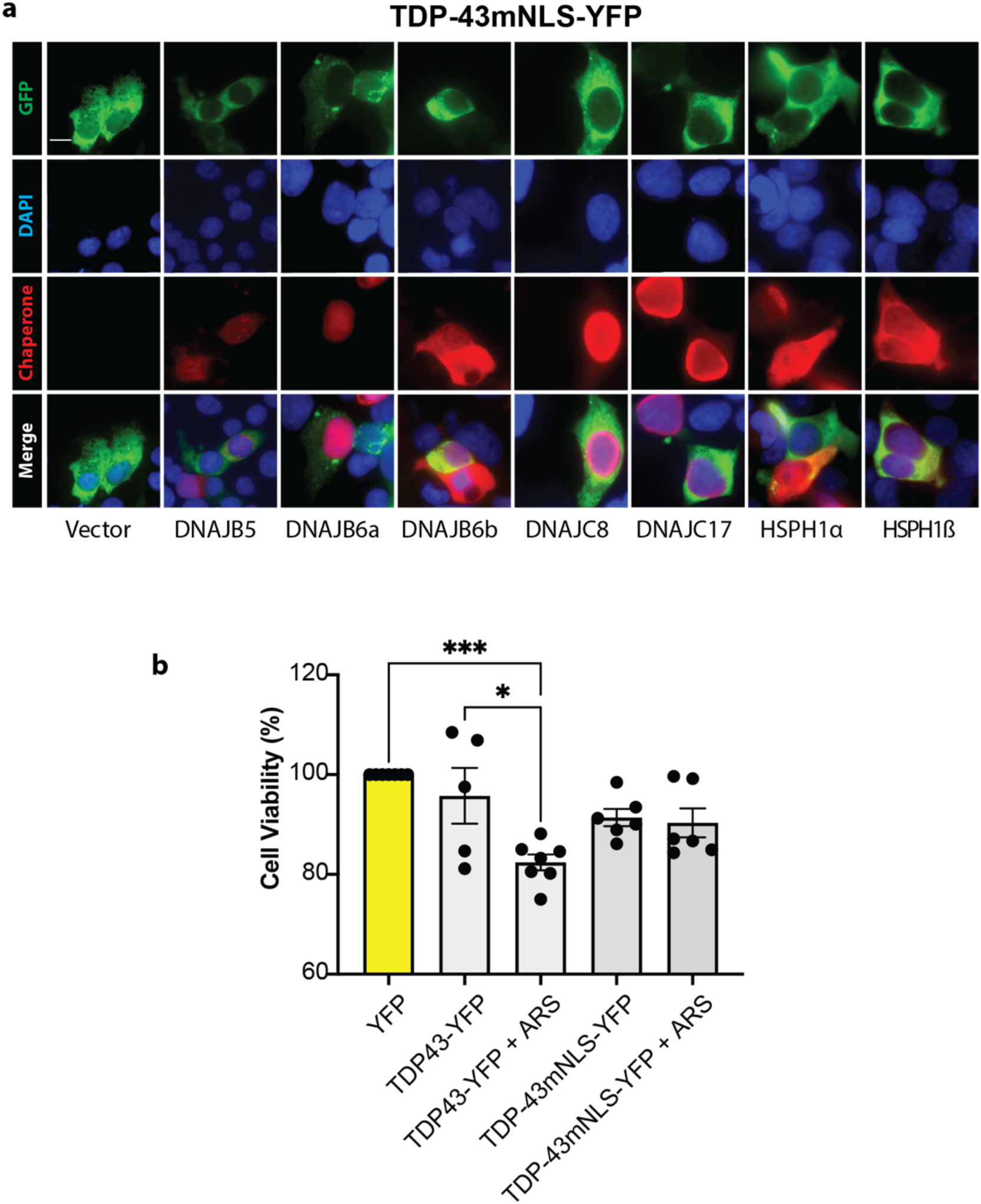
Expression of TDP-43-YFP, TDP-43mNLS-YFP, and chaperones in human cells. (**A**) Representative images showing expression of TDP-43mNLS-YFP and chaperones in HEK293 cells. Cell nuclei are stained with DAPI, and V5-tagged chaperones are detected by immunofluorescence. Scale bar, 20µm. (**B**) Cell viability for cells transfected with YFP, TDP-43-YFP, or TDP-43mNLS-YFP then treated with 5 µM sodium arsenite for 48 hours post transfection. Values are normalized to the YFP control in each experiment and represent mean ± SEM of 5-7 replicates. Statistical significance is determined by one-way ANOVA and Tukey’s test (*p < 0.05, ***p < 0.001).

**Supplementary Figure 10.**
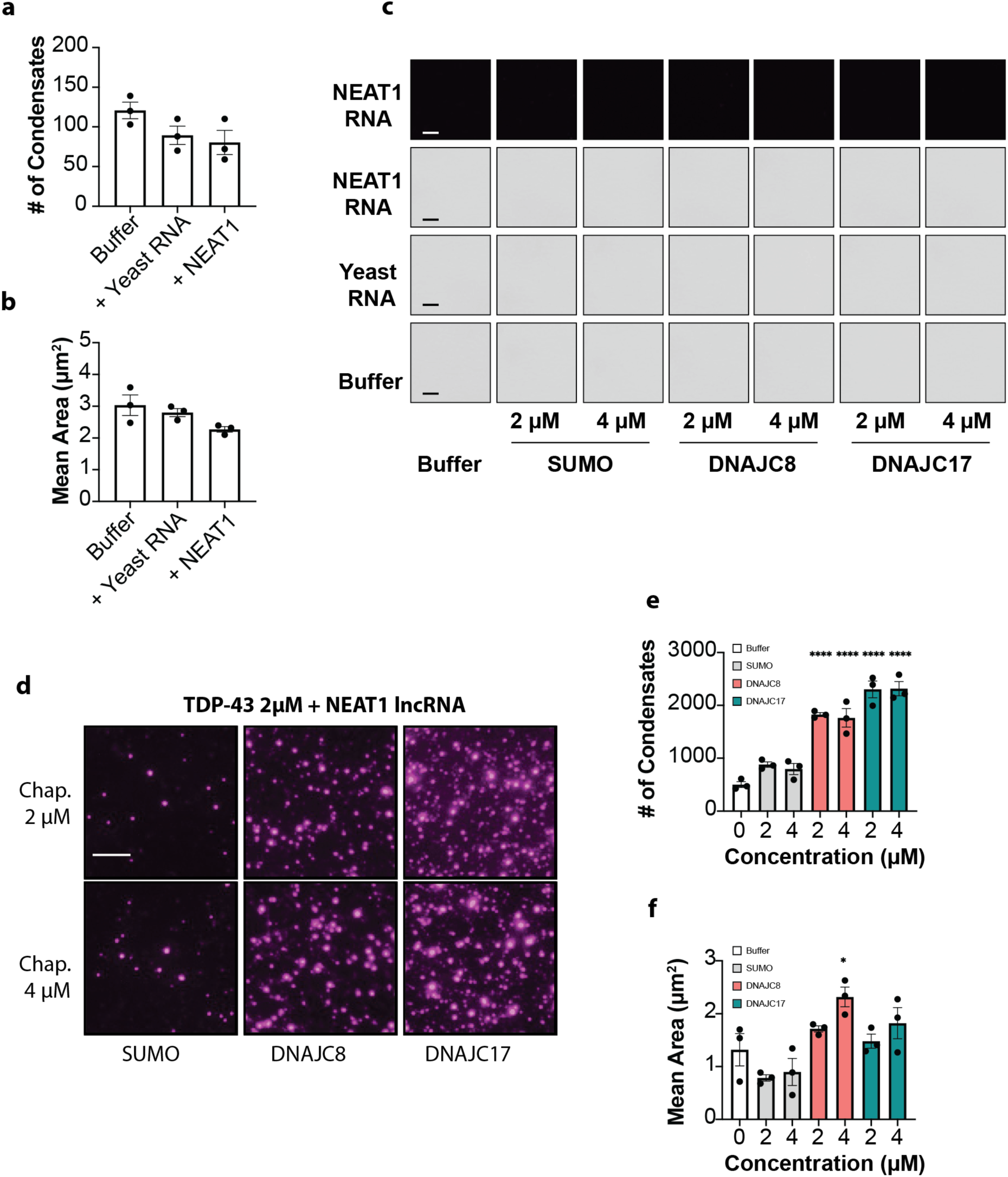
Additional controls confirming the enhancement of TDP-43 condensation in the presence of DNAJC8, DNAJC17, and RNA. (**A**) Quantification of TDP-43 condensate number in buffer with addition of yeast total RNA or NEAT1 lncRNA. Values represent mean ± SEM from three replicates. (**B**) Quantification of TDP-43 condensate area in buffer with addition of yeast total RNA or NEAT1 lncRNA. Values represent mean ± SEM from three replicates. (**C**) Control experiments performed in the absence of TDP-43 showing no detected condensates for the indicated RNAs and chaperones without TDP-43. Scale bar, 5 µm. (**D**) Representative images showing TDP-43 condensates in the presence of Cy5 labeled NEAT1 lncRNA in buffer with addition of SUMO, DNAJC8, or DNAJC17. Detection of condensates is through the Cy5 fluorescent label confirming the presence of NEAT1 lncRNA in the TDP-43 condensates. Scale bar, 5 µm. (**E, F**) Quantification of TDP-43 condensate number (E) or average area (F) in the presence of Cy5 labeled NEAT1 lncRNA in buffer with addition of SUMO, DNAJC8, or DNAJC17. Values represent mean ± SEM from three replicates. Statistical significance was determined relative to the buffer condition by one-way ANOVA and Dunnett’s multiple comparisons test (*p < 0.05, ****p < 0.0001).

## References

1. Mead, R.J., Shan, N., Reiser, H.J., Marshall, F. & Shaw, P.J. Amyotrophic lateral sclerosis: a neurodegenerative disorder poised for successful therapeutic translation. Nat Rev Drug Discov 22, 185–212 (2023).

2. Brown, D.G., Shorter, J. & Wobst, H.J. Emerging small-molecule therapeutic approaches for amyotrophic lateral sclerosis and frontotemporal dementia. Bioorg Med Chem Lett 30, 126942 (2020).

3. Wobst, H.J., Mack, K.L., Brown, D.G., Brandon, N.J. & Shorter, J. The clinical trial landscape in amyotrophic lateral sclerosis-Past, present, and future. Med Res Rev 40, 1352–1384 (2020).

4. Shemesh, N. et al. The landscape of molecular chaperones across human tissues reveals a layered architecture of core and variable chaperones. Nat Commun 12, 2180 (2021).

5. Portz, B., Lee, B.L. & Shorter, J. FUS and TDP-43 Phases in Health and Disease. Trends Biochem Sci 46, 550–563 (2021).

6. Ling, S.C., Polymenidou, M. & Cleveland, D.W. Converging mechanisms in ALS and FTD: disrupted RNA and protein homeostasis. Neuron 79, 416–438 (2013).

7. Meneses, A. et al. TDP-43 Pathology in Alzheimer’s Disease. Mol Neurodegener 16, 84 (2021).

8. Agra Almeida Quadros, A.R., et al. Cryptic splicing of stathmin-2 and UNC13A mRNAs is a pathological hallmark of TDP-43-associated Alzheimer’s disease. Acta Neuropathol 147, 9 (2024).

9. Nelson, P.T. et al. Limbic-predominant age-related TDP-43 encephalopathy (LATE): consensus working group report. Brain 142, 1503–1527 (2019).

10. McKee, A.C. et al. TDP-43 proteinopathy and motor neuron disease in chronic traumatic encephalopathy. J Neuropathol Exp Neurol 69, 918–929 (2010).

11. McKee, A.C., Stein, T.D., Kiernan, P.T. & Alvarez, V.E. The neuropathology of chronic traumatic encephalopathy. Brain Pathol 25, 350–364 (2015).

12. Ma, X.R. et al. TDP-43 represses cryptic exon inclusion in the FTD-ALS gene UNC13A. Nature 603, 124–130 (2022).

13. Brown, A.L. et al. TDP-43 loss and ALS-risk SNPs drive mis-splicing and depletion of UNC13A. Nature 603, 131–137 (2022).

14. Carey, J.L. & Guo, L. Liquid-Liquid Phase Separation of TDP-43 and FUS in Physiology and Pathology of Neurodegenerative Diseases. Front Mol Biosci 9, 826719 (2022).

15. Hallegger, M. et al. TDP-43 condensation properties specify its RNA-binding and regulatory repertoire. Cell 184, 4680–4696 e4622 (2021).

16. Grese, Z.R. et al. Specific RNA interactions promote TDP-43 multivalent phase separation and maintain liquid properties. EMBO Rep 22, e53632 (2021).

17. Huang, W.P. et al. Stress-induced TDP-43 nuclear condensation causes splicing loss of function and STMN2 depletion. Cell Rep 43, 114421 (2024).

18. Yu, H. et al. HSP70 chaperones RNA-free TDP-43 into anisotropic intranuclear liquid spherical shells. Science 371, eabb4309 (2021).

19. Modic, M. et al. Cross-Regulation between TDP-43 and Paraspeckles Promotes Pluripotency-Differentiation Transition. Mol Cell 74, 951–965 e913 (2019).

20. Fox, A.H., Nakagawa, S., Hirose, T. & Bond, C.S. Paraspeckles: Where Long Noncoding RNA Meets Phase Separation. Trends Biochem Sci 43, 124–135 (2018).

21. An, H., Williams, N.G. & Shelkovnikova, T.A. NEAT1 and paraspeckles in neurodegenerative diseases: A missing lnc found? Noncoding RNA Res 3, 243–252 (2018).

22. Fare, C.M. & Shorter, J. (Dis)Solving the problem of aberrant protein states. Dis Model Mech 14 (2021).

23. Khalil, B., Linsenmeier, M., Smith, C.L., Shorter, J. & Rossoll, W. Nuclear-import receptors as gatekeepers of pathological phase transitions in ALS/FTD. Mol Neurodegener 19, 8 (2024).

24. Meller, A. & Shalgi, R. The aging proteostasis decline: From nematode to human. Exp Cell Res 399, 112474 (2021).

25. Balch, W.E., Morimoto, R.I., Dillin, A. & Kelly, J.W. Adapting proteostasis for disease intervention. Science 319, 916–919 (2008).

26. Rosenzweig, R., Nillegoda, N.B., Mayer, M.P. & Bukau, B. The Hsp70 chaperone network. Nat Rev Mol Cell Biol 20, 665–680 (2019).

27. Kampinga, H.H. & Craig, E.A. The HSP70 chaperone machinery: J proteins as drivers of functional specificity. Nat Rev Mol Cell Biol 11, 579–592 (2010).

28. Mashaghi, A. et al. Alternative modes of client binding enable functional plasticity of Hsp70. Nature 539, 448–451 (2016).

29. Zhang, R., Malinverni, D., Cyr, D.M., Rios, P.D.L. & Nillegoda, N.B. J-domain protein chaperone circuits in proteostasis and disease. Trends in Cell Biology 33, 30–47 (2023).

30. Piette, B.L. et al. Comprehensive interactome profiling of the human Hsp70 network highlights functional differentiation of J domains. Mol Cell 81, 2549–2565 e2548 (2021).

31. Cvitkovic, I. & Jurica, M.S. Spliceosome database: a tool for tracking components of the spliceosome. Nucleic Acids Res 41, D132–141 (2013).

32. Pascarella, A. et al. DNAJC17 is localized in nuclear speckles and interacts with splicing machinery components. Sci Rep 8, 7794 (2018).

33. Allegakoen, D.V., Kwong, K., Morales, J., Bivona, T.G. & Sabnis, A.J. The essential chaperone DNAJC17 activates HSP70 to coordinate RNA splicing and G2-M progression. bioRxiv, 2023.2010.2025.564066 (2023).

34. Ajit Tamadaddi, C. & Sahi, C. J domain independent functions of J proteins. Cell Stress Chaperones 21, 563–570 (2016).

35. Mattoo, R.U.H., Sharma, S.K., Priya, S., Finka, A. & Goloubinoff, P. Hsp110 is a bona fide chaperone using ATP to unfold stable misfolded polypeptides and reciprocally collaborate with Hsp70 to solubilize protein aggregates. J Biol Chem 288, 21399–21411 (2013).

36. Gillis, J. et al. The DNAJB6 and DNAJB8 protein chaperones prevent intracellular aggregation of polyglutamine peptides. J Biol Chem 288, 17225–17237 (2013).

37. Oh, H.J., Chen, X. & Subjeck, J.R. Hsp110 protects heat-denatured proteins and confers cellular thermoresistance. J Biol Chem 272, 31636–31640 (1997).

38. Hu, L. et al. A first-in-class inhibitor of Hsp110 molecular chaperones of pathogenic fungi. Nat Commun 14, 2745 (2023).

39. Johnson, B.S., McCaffery, J.M., Lindquist, S. & Gitler, A.D. A yeast TDP-43 proteinopathy model: Exploring the molecular determinants of TDP-43 aggregation and cellular toxicity. Proc Natl Acad Sci U S A 105, 6439–6444 (2008).

40. Johnson, B.S. et al. TDP-43 is intrinsically aggregation-prone, and amyotrophic lateral sclerosis-linked mutations accelerate aggregation and increase toxicity. J Biol Chem 284, 20329–20339 (2009).

41. Jackrel, M.E. et al. Potentiated Hsp104 Variants Antagonize Diverse Proteotoxic Misfolding Events. Cell 156, 170–182 (2014).

42. Becker, L.A. et al. Therapeutic reduction of ataxin-2 extends lifespan and reduces pathology in TDP-43 mice. Nature 544, 367–371 (2017).

43. Elden, A.C. et al. Ataxin-2 intermediate-length polyglutamine expansions are associated with increased risk for ALS. Nature 466, 1069–1075 (2010).

44. Kim, H.J. et al. Therapeutic modulation of eIF2alpha phosphorylation rescues TDP-43 toxicity in amyotrophic lateral sclerosis disease models. Nat Genet 46, 152–160 (2014).

45. Armakola, M. et al. Inhibition of RNA lariat debranching enzyme suppresses TDP-43 toxicity in ALS disease models. Nat Genet 44, 1302–1309 (2012).

46. Tardiff, D.F., Tucci, M.L., Caldwell, K.A., Caldwell, G.A. & Lindquist, S. Different 8-hydroxyquinolines protect models of TDP-43 protein, alpha-synuclein, and polyglutamine proteotoxicity through distinct mechanisms. J Biol Chem 287, 4107–4120 (2012).

47. Lin, J. et al. Design principles to tailor Hsp104 therapeutics. Cell Rep 43, 115005 (2024).

48. Shaner, N.C. et al. A bright monomeric green fluorescent protein derived from Branchiostoma lanceolatum. Nat Methods 10, 407–409 (2013).

49. Gitcho, M.A. et al. TARDBP 3’-UTR variant in autopsy-confirmed frontotemporal lobar degeneration with TDP-43 proteinopathy. Acta Neuropathol 118, 633–645 (2009).

50. Gasset-Rosa, F. et al. Cytoplasmic TDP-43 De-mixing Independent of Stress Granules Drives Inhibition of Nuclear Import, Loss of Nuclear TDP-43, and Cell Death. Neuron 102, 339–357 e337 (2019).

51. Polymenidou, M. et al. Long pre-mRNA depletion and RNA missplicing contribute to neuronal vulnerability from loss of TDP-43. Nat Neurosci 14, 459–468 (2011).

52. Bobkova, N.V. et al. Exogenous Hsp70 delays senescence and improves cognitive function in aging mice. Proc Natl Acad Sci U S A 112, 16006–16011 (2015).

53. Yang, S., Huang, S., Gaertig, M.A., Li, X.J. & Li, S. Age-dependent decrease in chaperone activity impairs MANF expression, leading to Purkinje cell degeneration in inducible SCA17 mice. Neuron 81, 349–365 (2014).

54. Santra, M., Dill, K.A. & de Graff, A.M.R. Proteostasis collapse is a driver of cell aging and death. Proc Natl Acad Sci U S A 116, 22173–22178 (2019).

55. Shpund, S. & Gershon, D. Alterations in the chaperone activity of HSP70 in aging organisms. Arch Gerontol Geriatr 24, 125–131 (1997).

56. Malinverni, D. et al. Data-driven large-scale genomic analysis reveals an intricate phylogenetic and functional landscape in J-domain proteins. Proc Natl Acad Sci U S A 120, e2218217120 (2023).

57. Rebeaud, M.E., Mallik, S., Goloubinoff, P. & Tawfik, D.S. On the evolution of chaperones and cochaperones and the expansion of proteomes across the Tree of Life. Proc Natl Acad Sci U S A 118, e2020885118 (2021).

58. Farhan, S.M.K. et al. Exome sequencing in amyotrophic lateral sclerosis implicates a novel gene, DNAJC7, encoding a heat-shock protein. Nat Neurosci 22, 1966–1974 (2019).

59. Fleming, A.C., Rao, N.R., Wright, M., Savas, J.N. & Kiskinis, E. The ALS-associated co-chaperone DNAJC7 mediates neuroprotection against proteotoxic stress by modulating HSF1 activity. eLife (2025).

60. Park, S.K. et al. Overexpression of the essential Sis1 chaperone reduces TDP-43 effects on toxicity and proteolysis. PLoS Genet 13, e1006805 (2017).

61. Carrasco, J. et al. Metamorphism in TDP-43 prion-like domain determines chaperone recognition. Nat Commun 14, 466 (2023).

62. San Gil, R., et al. A transient protein folding response targets aggregation in the early phase of TDP-43-mediated neurodegeneration. Nat Commun 15, 1508 (2024).

63. Abayev-Avraham, M., Salzberg, Y., Gliksberg, D., Oren-Suissa, M. & Rosenzweig, R. DNAJB6 mutants display toxic gain of function through unregulated interaction with Hsp70 chaperones. Nat Commun 14, 7066 (2023).

64. Resnick, S.J. et al. A multiplex platform to identify mechanisms and modulators of proteotoxicity in neurodegeneration. bioRxiv, 2022.2009.2019.508444 (2022).

65. Stein, K.C., Bengoechea, R., Harms, M.B., Weihl, C.C. & True, H.L. Myopathy-causing mutations in an HSP40 chaperone disrupt processing of specific client conformers. J Biol Chem 289, 21120–21130 (2014).

66. Bengoechea, R. et al. P.5.11 LGMD1D mutations in DNAJB6 disrupt disaggregation of TDP-43. Neuromuscular Disorders 23, 767 (2013).

67. Francois-Moutal, L. et al. Heat shock protein Grp78/BiP/HspA5 binds directly to TDP-43 and mitigates toxicity associated with disease pathology. Sci Rep 12, 8140 (2022).

68. Ormeno, F. et al. Chaperone Mediated Autophagy Degrades TDP-43 Protein and Is Affected by TDP-43 Aggregation. Front Mol Neurosci 13, 19 (2020).

69. Pinarbasi, E.S. et al. Active nuclear import and passive nuclear export are the primary determinants of TDP-43 localization. Sci Rep 8, 7083 (2018).

70. Charoniti, E. et al. TARDBP p.I383V, a recurrent alteration in Greek FTD patients. J Neurol Sci 428, 117566 (2021).

71. Agrawal, S., Jain, M., Yang, W.Z. & Yuan, H.S. Frontotemporal dementia-linked P112H mutation of TDP-43 induces protein structural change and impairs its RNA binding function. Protein Sci 30, 350–365 (2021).

72. Harrison, A.F. & Shorter, J. RNA-binding proteins with prion-like domains in health and disease. Biochem J 474, 1417–1438 (2017).

73. Chen, H.J. et al. RRM adjacent TARDBP mutations disrupt RNA binding and enhance TDP-43 proteinopathy. Brain 142, 3753–3770 (2019).

74. Ling, S.C. et al. ALS-associated mutations in TDP-43 increase its stability and promote TDP-43 complexes with FUS/TLS. Proc Natl Acad Sci U S A 107, 13318–13323 (2010).

75. Arseni, D. et al. TDP-43 forms amyloid filaments with a distinct fold in type A FTLD-TDP. Nature 620, 898–903 (2023).

76. Arseni, D. et al. Structure of pathological TDP-43 filaments from ALS with FTLD. Nature 601, 139–143 (2022).

77. Conicella, A.E. et al. TDP-43 alpha-helical structure tunes liquid-liquid phase separation and function. Proc Natl Acad Sci U S A 117, 5883–5894 (2020).

78. Conicella, A.E., Zerze, G.H., Mittal, J. & Fawzi, N.L. ALS Mutations Disrupt Phase Separation Mediated by alpha-Helical Structure in the TDP-43 Low-Complexity C-Terminal Domain. Structure 24, 1537–1549 (2016).

79. Yan, X. et al. Intra-condensate demixing of TDP-43 inside stress granules generates pathological aggregates. bioRxiv, 2024.2001.2023.576837 (2024).

80. Copley, K.E. et al. Short RNA chaperones promote aggregation-resistant TDP-43 conformers to mitigate neurodegeneration. bioRxiv, 2024.2012.2014.628507 (2024).

81. Lang, R., Hodgson, R.E. & Shelkovnikova, T.A. TDP-43 in nuclear condensates: where, how, and why. Biochemical Society Transactions 52, 1809–1825 (2024).

82. Bolognesi, B. et al. The mutational landscape of a prion-like domain. Nat Commun 10, 4162 (2019).

83. Johnson, O.T., Nadel, C.M., Carroll, E.C., Arhar, T. & Gestwicki, J.E. Two distinct classes of cochaperones compete for the EEVD motif in heat shock protein 70 to tune its chaperone activities. J Biol Chem 298, 101697 (2022).

84. Dragovic, Z., Broadley, S.A., Shomura, Y., Bracher, A. & Hartl, F.U. Molecular chaperones of the Hsp110 family act as nucleotide exchange factors of Hsp70s. EMBO J 25, 2519–2528 (2006).

85. Shaner, L., Wegele, H., Buchner, J. & Morano, K.A. The yeast Hsp110 Sse1 functionally interacts with the Hsp70 chaperones Ssa and Ssb. J Biol Chem 280, 41262–41269 (2005).

86. Shaner, L., Trott, A., Goeckeler, J.L., Brodsky, J.L. & Morano, K.A. The function of the yeast molecular chaperone Sse1 is mechanistically distinct from the closely related hsp70 family. J Biol Chem 279, 21992–22001 (2004).

87. Saxena, A. et al. Human heat shock protein 105/110 kDa (Hsp105/110) regulates biogenesis and quality control of misfolded cystic fibrosis transmembrane conductance regulator at multiple levels. J Biol Chem 287, 19158–19170 (2012).

88. Seike, T. et al. Site-specific photo-crosslinking of Hsc70 with the KFERQ pentapeptide motif in a chaperone-mediated autophagy and microautophagy substrate in mammalian cells. Biochem Biophys Res Commun 736, 150515 (2024).

89. Huang, S.P., Tsai, M.Y., Tzou, Y.M., Wu, W.G. & Wang, C. Aspartyl residue 10 is essential for ATPase activity of rat hsc70. J Biol Chem 268, 2063–2068 (1993).

90. Saito, Y., Yamagishi, N. & Hatayama, T. Different localization of Hsp105 family proteins in mammalian cells. Exp Cell Res 313, 3707–3717 (2007).

91. Pokhrel, S., Devi, S. & Gestwicki, J.E. Chaperone-dependent and chaperone-independent functions of carboxylate clamp tetratricopeptide repeat (CC-TPR) proteins. Trends Biochem Sci 50, 121–133 (2025).

92. Faust, O. et al. HSP40 proteins use class-specific regulation to drive HSP70 functional diversity. Nature 587, 489–494 (2020).

93. Shorter, J. & Southworth, D.R. Spiraling in Control: Structures and Mechanisms of the Hsp104 Disaggregase. Cold Spring Harb Perspect Biol 11 (2019).

94. Sweeny, E.A. & Shorter, J. Mechanistic and Structural Insights into the Prion-Disaggregase Activity of Hsp104. J Mol Biol 428, 1870–1885 (2016).

95. Erives, A.J. & Fassler, J.S. Metabolic and chaperone gene loss marks the origin of animals: evidence for Hsp104 and Hsp78 chaperones sharing mitochondrial enzymes as clients. PLoS One 10, e0117192 (2015).

96. Kabani, M. & Martineau, C.N. Multiple hsp70 isoforms in the eukaryotic cytosol: mere redundancy or functional specificity? Curr Genomics 9, 338–248 (2008).

97. Werner-Washburne, M., Stone, D.E. & Craig, E.A. Complex interactions among members of an essential subfamily of hsp70 genes in Saccharomyces cerevisiae. Mol Cell Biol 7, 2568–2577 (1987).

98. Hasin, N., Cusack, S.A., Ali, S.S., Fitzpatrick, D.A. & Jones, G.W. Global transcript and phenotypic analysis of yeast cells expressing Ssa1, Ssa2, Ssa3 or Ssa4 as sole source of cytosolic Hsp70-Ssa chaperone activity. BMC Genomics 15, 194 (2014).

99. Sanchez, Y. et al. Genetic evidence for a functional relationship between Hsp104 and Hsp70. J Bacteriol 175, 6484–6491 (1993).

100. Uhrigshardt, H., Singh, A., Kovtunovych, G., Ghosh, M. & Rouault, T.A. Characterization of the human HSC20, an unusual DnaJ type III protein, involved in iron-sulfur cluster biogenesis. Hum Mol Genet 19, 3816–3834 (2010).

101. Voisine, C. et al. Jac1, a mitochondrial J-type chaperone, is involved in the biogenesis of Fe/S clusters in Saccharomyces cerevisiae. Proc Natl Acad Sci U S A 98, 1483–1488 (2001).

102. Shapiro, O. et al. Assays to measure small molecule Hsp70 agonist activity in vitro and in vivo. Anal Biochem 697, 115712 (2025).

103. Chuang, E. et al. Defining a small-molecule stimulator of the human Hsp70-disaggregase system with selectivity for DnaJB proteins. bioRxiv, 2024.2001.2011.575109 (2024).

104. Zhang, J.Z. et al. De novo designed Hsp70 activator dissolves intracellular condensates. Cell Chem Biol 32, 463–473 e466 (2025).

105. Osterlund, N. et al. The C-terminal domain of the antiamyloid chaperone DNAJB6 binds to amyloid-beta peptide fibrils and inhibits secondary nucleation. J Biol Chem 299, 105317 (2023).

106. Bai, S. et al. DNAJB7 is dispensable for male fertility in mice. Reprod Biol Endocrinol 21, 32 (2023).

107. Scurr, M.J. et al. Cancer Antigen Discovery Is Enabled by RNA Sequencing of Highly Purified Malignant and Nonmalignant Cells. Clin Cancer Res 26, 3360–3370 (2020).

108. Uhlén, M. et al. Tissue-based map of the human proteome. Science 347, 1260419 (2015).

109. Ryder, B.D. et al. DNAJB8 oligomerization is mediated by an aromatic-rich motif that is dispensable for substrate activity. Structure 32, 662–678 e668 (2024).

110. Zhong, X.Y., Ding, J.H., Adams, J.A., Ghosh, G. & Fu, X.D. Regulation of SR protein phosphorylation and alternative splicing by modulating kinetic interactions of SRPK1 with molecular chaperones. Genes Dev 23, 482–495 (2009).

111. Ito, N. et al. A novel nuclear DnaJ protein, DNAJC8, can suppress the formation of spinocerebellar ataxia 3 polyglutamine aggregation in a J-domain independent manner. Biochem Biophys Res Commun 474, 626–633 (2016).

112. Raut, S. et al. Co-evolution of spliceosomal disassembly interologs: crowning J-protein component with moonlighting RNA-binding activity. Curr Genet 65, 561–573 (2019).

113. Sahi, C., Lee, T., Inada, M., Pleiss, J.A. & Craig, E.A. Cwc23, an essential J protein critical for pre-mRNA splicing with a dispensable J domain. Mol Cell Biol 30, 33–42 (2010).

114. Suk, T.R. & Rousseaux, M.W.C. The role of TDP-43 mislocalization in amyotrophic lateral sclerosis. Mol Neurodegener 15, 45 (2020).

115. McGurk, L. et al. Poly(ADP-Ribose) Prevents Pathological Phase Separation of TDP-43 by Promoting Liquid Demixing and Stress Granule Localization. Mol Cell 71, 703–717 e709 (2018).

116. Lang, R., Hodgson, R.E. & Shelkovnikova, T.A. TDP-43 in nuclear condensates: where, how, and why. Biochem Soc Trans 52, 1809–1825 (2024).

117. Shen, Y. & Hendershot, L.M. ERdj3, a stress-inducible endoplasmic reticulum DnaJ homologue, serves as a cofactor for BiP’s interactions with unfolded substrates. Mol Biol Cell 16, 40–50 (2005).

118. Pathak, D., Berthet, A. & Nakamura, K. Energy failure: does it contribute to neurodegeneration? Ann Neurol 74, 506–516 (2013).

119. Guillaud, L. et al. Loss of intracellular ATP affects axoplasmic viscosity and pathological protein aggregation in mammalian neurons. Sci Adv 11, eadq6077 (2025).

120. Amado, D.A. & Davidson, B.L. Gene therapy for ALS: A review. Mol Ther 29, 3345–3358 (2021).

121. Perera, A., Brock, O., Ahmed, A., Shaw, C. & Ashkan, K. Taking the knife to neurodegeneration: a review of surgical gene therapy delivery to the CNS. Acta Neurochir (Wien) 166, 136 (2024).

122. Challis, R.C. et al. Adeno-Associated Virus Toolkit to Target Diverse Brain Cells. Annu Rev Neurosci 45, 447–469 (2022).

123. Tervo, D.G. et al. A Designer AAV Variant Permits Efficient Retrograde Access to Projection Neurons. Neuron 92, 372–382 (2016).

124. Wilkins, O.G. et al. Creation of de novo cryptic splicing for ALS and FTD precision medicine. Science 386, 61–69 (2024).

125. Gietz, R.D. & Schiestl, R.H. High-efficiency yeast transformation using the LiAc/SS carrier DNA/PEG method. Nat Protoc 2, 31–34 (2007).

126. Jumper, J. et al. Highly accurate protein structure prediction with AlphaFold. Nature 596, 583–589 (2021).

